# Dissecting the Cellular Genetics of Cardiovascular Disease Through Endothelial and Immune Compartments Profiling

**DOI:** 10.1101/2025.11.10.687750

**Authors:** Stephen Watt, Matiss Ozols, Arianna Landini, Bin Liu, Sodbo Sharapov, Daniela Zanotti, Gabriela Ivanova, Charles Solomon, Xu Dong Yang, Tolulope Balogun, Nicole Staudt, Wei Li, Fatin Al-Janabi, Tom R. Webb, David G. McVey, Nicholas Morrell, Martin Bennett, Nilesh J. Samani, Stefan Gräf, Shu Ye, Nicola Pirastu, Nicole Soranzo, Mattia Frontini

**Affiliations:** Wellcome Sanger Institute, Wellcome Genome Campus, Hixton, UK; Cell Matrix Biology & Regenerative Medicine, Faculty of Biology, Medicine and Health, The University of Manchester, Manchester, UK; Human Technopole, Fondazione Human Technopole, Milan, Italy; Victor Phillip Dahdaleh Heart & Lung Research Institute, University of Cambridge, Cambridge, UK; Politecnico di Milano, Milano, Italy; Division of Cardiovascular Sciences, Leicester British Heart Foundation Centre of Research Excellence and NIHR Leicester Biomedical Research Centre, University of Leicester, Leicester, UK; Department of Clinical and Biomedical Sciences, University of Exeter, Exeter, UK; National University of Singapore, Singapore; First affiliated hospital of Shantou University medical college, Shantou, CN

## Abstract

**Background:** Non-communicable diseases such as coronary artery disease, atrial fibrillation, type 2 diabetes, hypertension, and others share endothelial dysfunction as one of their underlying features. The endothelium, as the interface between blood and vasculature, shapes disease onset and progression through its response to environmental cues. However, while the genetic component of these diseases has been captured by genome wide association studies (GWAS), which also highlighted a shared immune component, it remains unclear which of these disease loci exerts their effects through endothelial cells. This study identifies, and quantifies, the genetic determinants of endothelial cells molecular traits and their overlap to the common genetic variation component of these diseases.

**Methods:** We generated genotype, RNA-sequencing, H3K27ac ChIP-sequencing, ATAC-sequencing, and endothelial cells barrier stimuli response measurements for 100 samples of human umbilical vein endothelial cells. These were used to identify quantitative trait loci (QTL) for gene expression, transcriptional isoform usage, splice junction usage, chromatin activity and barrier response. We applied statistical colocalisation to identify the overlap between data layers, and to explain molecular QTLs contribution to GWAS disease loci.

**Results:** We used molecular QTLs to identify the regulatory features of 8,214 genes, representing 36% of all expressed genes in endothelial cells. We also identified the molecular mechanisms underlying 815 loci across 16 disease GWAS. These represent between 29% and 40% of all loci for each disease, compared to the previous average of 23%. This is due to the choice of a cell type often underrepresented in tissue level data, and the inclusion of isoform, splicing and chromatin activity datasets. Furthermore, we compared the endothelial cells molecular QTLs with similar datasets in monocytes, neutrophils and CD4 T lymphocytes to shed light on the interplay between the endothelial and the immune compartments in these diseases. We identified loci acting through both the endothelial and the immune compartment, mostly with the same directionality of effect, and endothelial specific ones.

**Conclusions:** This work expands the knowledge of the mechanisms and genes underlying the effect of common genetic variation on non-communicable diseases having endothelial dysfunction as a shared feature. It also illustrates the interplay between endothelial cells and immune cell types in these diseases, highlighting shared and unique pathways.

## Introduction

Endothelial cells (ECs) form a single cell layer lining blood and lymphatic vessels, ensuring proper vascular function (Krüger-Genge et al. 2019). They play critical roles in (i) maintaining haemostasis (Yau, Teoh, and Verma 2015); (ii) regulating vascular tone (Baumgartner-Parzer and Waldhäusl 2001); (iii) controlling the trafficking of immune cells, nutrients and other substances between blood circulation and tissues (Claesson-Welsh, Dejana, and McDonald 2021). Exposure to cholesterol, free radicals, and other factors impairs, or disrupts, these functions, leading to endothelial dysfunction (Rajendran et al. 2013), Characterized by reduced ability to vasodilate, resulting in hypertension, a shift towards a prothrombotic and proinflammatory state, and vascular leakage (Rajendran et al. 2013). These changes promote atherosclerotic plaque formation, and leukocyte tissue infiltration. Endothelial dysfunction also precedes insulin resistance and type 2 diabetes mellitus contributing to its complications (Hadi and Suwaidi 2007), and it is a predictor of cardiovascular disease (Gimbrone and García-Cardeña 2016).

While endothelial dysfunction has a genetic component, identified by familial studies and monogenic disorders (Jones and Hingorani 2005), this has never been explored in depth. Meanwhile studies of genetic risk loci for complex diseases characterised by endothelial dysfunction have identified thousands of contributing variants (Keaton et al. 2024) (Dönertaş et al. 2021) (Zheng et al. 2021) (Nielsen et al. 2018) (Mahajan et al. 2022) (Mishra et al. 2022) (Aragam et al. 2022), mostly thought to affect disease through gene regulation effects (Maurano et al. 2012). However, the tissues and mechanisms underlying these associations remain largely unknown, with only 22 coronary artery disease (CAD) loci attributed to endothelial cells (Aragam et al. 2022).

Recent studies have deepened our understanding of the regulatory landscape underlying CAD by integrating GWAS with molecular QTL from arterial tissues (Liu et al. 2018) (Stolze et al. 2020) (Solomon et al. 2022) (Aherrahrou et al. 2023). Single-cell analyses further highlighted that CAD associated variants are enriched in open chromatin regions of endothelial and vascular smooth muscle cells (VSMCs) (Turner et al. 2022) (Örd et al. 2021) (Adelus et al. 2024). CRISPR-based perturbation studies have demonstrated that candidate causal genes and variants can directly modulate endothelial dysfunction (Wünnemann et al. 2023). Together, these findings support a CAD risk model mediated through cellular dysfunction within the vascular wall. Nevertheless, a systematic comparison of the regulatory architecture across endothelial and immune cell types is still lacking.

Human umbilical vein endothelial cells (HUVECs) are a model for studying endothelial biology, given their accessibility, similarity to vascular endothelial cells, and relevance in both physiological and pathological contexts (Onat et al. 2011). To identify disease-associated variants acting through endothelial cells and to determine the contribution of genetic variation to their regulatory landscape, we generated a new multi-omics dataset composed of 100 HUVEC samples capturing multiple layers of gene regulation, including gene and transcript expression, and chromatin activity and occupancy.

## Results

### A new multi-omic atlas of human endothelial cells

HUVECs were harvested, alongside VSMCs, from umbilical cords provided by the Anthony Nolan Foundation and cryopreserved (Solomon et al. 2022). A total of 100 samples were retrieved, grown, and at passage 2 divided to harvest RNA for RNA-sequencing (RNA-seq; n=100); formaldehyde fixed chromatin for histone 3 lysine 27 acetylation chromatin immunoprecipitation (H3K27ac ChIP-seq; n=98); cells for assay for transposase-accessible chromatin (ATAC-seq; n=100; **Fig. 1A, Table S1**) and two wells (on separate arrays) for functional assays by Electric Cell-substrate Impedance Sensing (ECIS; **Methods**). RNA-seq was used to quantify gene expression, transcriptional isoforms and splice junction usage (Chen et al. 2016). H3K27ac ChIP-seq was used to identify and quantify promoters and transcriptional enhancers (Chen et al. 2016)), while ATAC-seq was used to map regions of open chromatin with regulatory potential (Buenrostro et al. 2013).

**Fig. 1.**
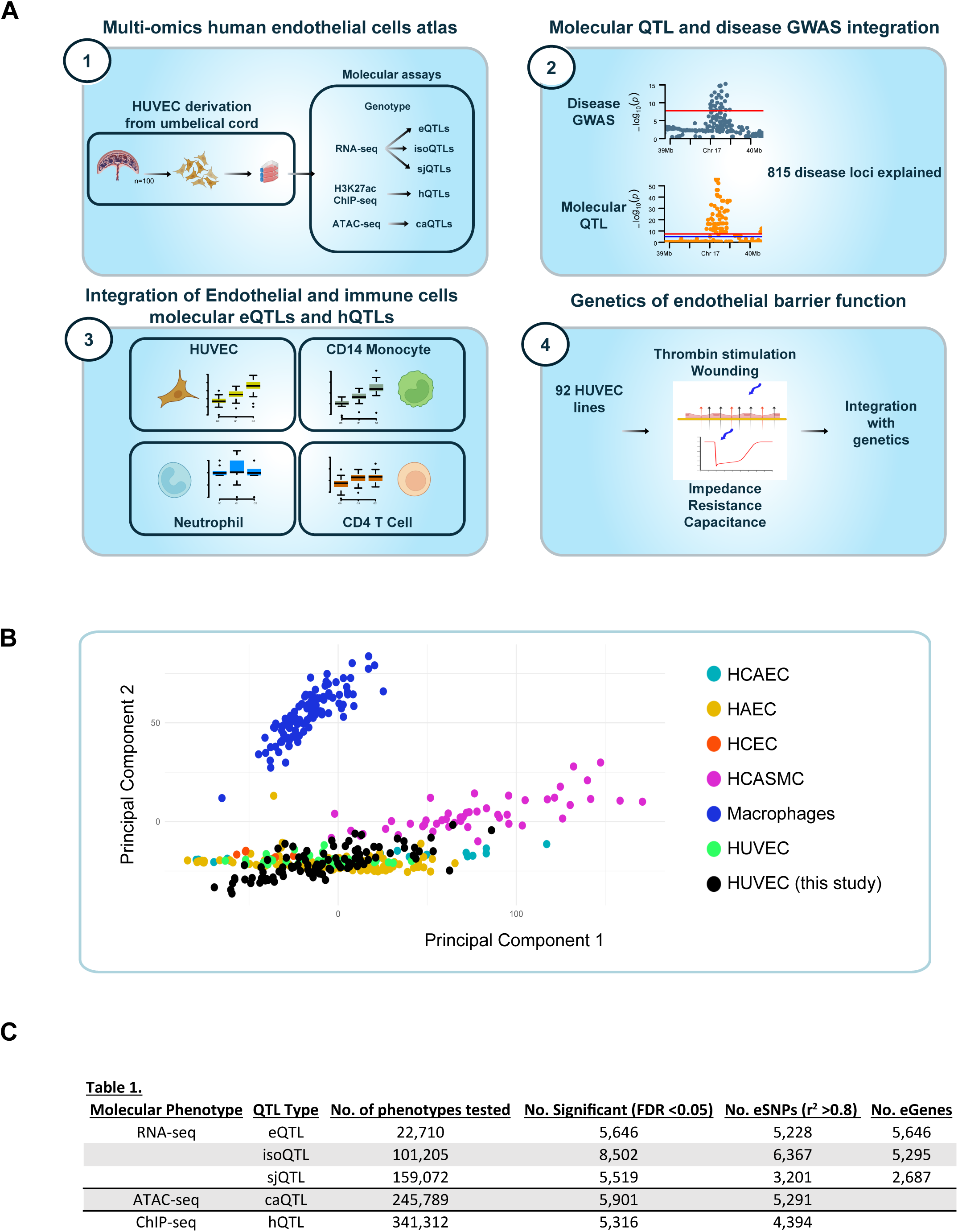
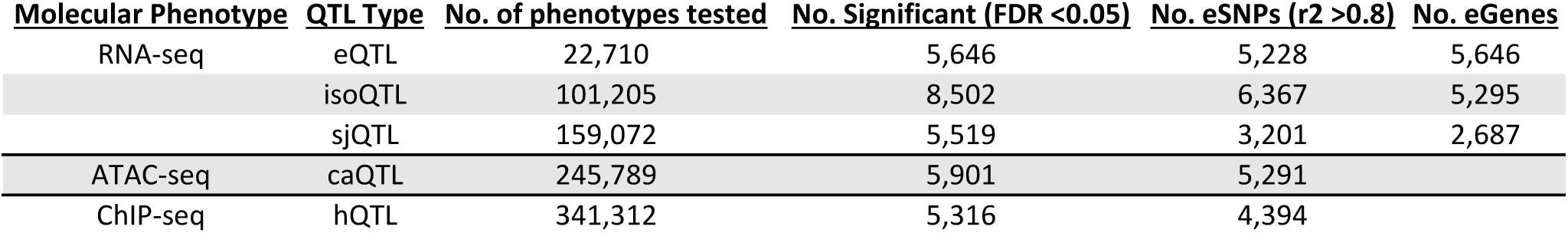
Study design. **A.** HUVEC were harvested from 100 placenta samples. Cells were expanded for two passages and used to generate the following dataset: genotype, RNA-seq, H3K27ac ChIP-seq, ATAC-seq, endothelial barrier formation, wounding assay, and thrombin activation. The dataset were used to identify the following associations with genotype: gene expression (eQTLs), mRNA isoforms usage (isoQTLs), splice junctions usage (sQTLs), H3K27ac peaks (hQTLs), ATAC-seq peaks (caQTLs) and the different cell functional phenotypes. These were then used in statistical colocalization with loci associated with diseases with underlying endothelial dysfunction GWAS. **B. Principal components analysis of gene expression summary statistics of endothelial cells from different vascular beds.** Scatterplot of the first (PC1) versus the second (PC2) components of gene expression data for different types of endothelial cells (HCAEC, human corneal endothelial cells; HAEC, human arterial endothelial cells; HCEC, human coronary artery endothelial cells; HUVEC, human umbilical vein endothelial cells) and other related cell types (HCASMC, human coronary artery smooth muscle cells and macrophages; **Suppl. Table S2** for data sources). **C.** Table displaying the number of significant molecular QTLs per trait tested.

Endothelial cells from different vascular beds exhibit subtle phenotypic differences (Z. Zhao et al. 2015; Gifre-Renom et al. 2022). To assess how our samples relate to other endothelial populations, we performed principal component analysis for gene expression including coronary aorta endothelial cells (HCAEC), aortic endothelial cells (HAEC), corneal endothelial cells (HCEC) and HUVECs from additional sources. The analysis revealed substantial overlap among most endothelial types, with HCEC displaying moderate divergence from the others (**Fig. 1B**; Hotelling’s T^2^ permutation test; **Table S2**). Pairwise differential gene expression (DGE) analysis, followed by Gene Ontology (GO) enrichment, highlighted pathways related to transmembrane signalling, CXCR chemokine binding, cell adhesion and inflammatory response ( **Table S3**).

To assess the impact of genetic variation on gene expression, we tested 22,710 expressed genes (non-zero expression in ≥ 20% of donors; **Methods**) using TensorQTL (Taylor-Weiner et al. 2019) and identified **5,646** eQTLs at a global FDR < 0.05 (**Methods; Table S4, Fig. 1C**).

We compared HUVEC eQTLs with those identified in HAEC (n=4,911 eQTLs) generated utilising both RNA-seq (53 donors) and microarray data (150 donors) (Romanoski et al. 2010) (Stolze et al. 2020). We identified **1,018** eQTLs with matched lead variants (LD r^2^≥0.8), associated with **1,834** distinct genes (**Suppl. Fig. S1A; Table S5**). GO enrichment for eQTL genes unique to each dataset retrieved terms associated with cell-cell adhesion (*p*=6.77×10^−6^) HUVEC specific and vasculature (*p*=1.73×10^−9^) and heart development (*p*=3.14×10^−6^), HAEC specific. These findings suggest that HUVEC- and HAEC-derived eQTLs capture complementary biological processes.

Alternative splicing represents an independent layer of gene regulation (Park et al. 2018). We quantified how genetic variation affects it using two approaches: reference-based annotation, Salmon (Patro et al. 2017) to determine isoform usage, and reference-agnostic annotation, Leafcutter (Li et al. 2018) to map splice junctions. These were then used to determine isoform usage (isoQTL) and splice junction (sjQTL) QTLs, respectively. We identified **8,502** isoQTLs (**5,295** genes), and **5,519** sjQTLs (**2,687** genes; 5% FDR; **Fig. 1C, Suppl. Table S4**). Active chromatin enables functional fine-mapping and mechanistic interpretation (Farh et al. 2015). To this end, we generated H3K27ac ChIP-seq (n=98) and chromatin occupancy ATAC-seq (n=100) profiles. Data quality was assessed from relevant metrics (e.g. fraction of reads in peaks, **Methods**) and visual inspection (**Suppl. Fig. S1B**, **Methods**). For each assay, we constructed a union peak set, identifying ∼350,000 H3K27ac peaks and ∼240,000 ATAC peaks (**Suppl. Fig. S1C-D**). Normalized read counts at peak regions were then used (**Methods**), yielding **5,316** hQTLs and **5,901** caQTLs at a global false discovery rate [gFDR] < 0.05 (**Fig. 1C, Table S4**). In total, hQTLs and caQTLs represented **8,541** genomic intervals, with a 29% overlap.

### Cross-layer QTL colocalisation reveals genetic architecture of gene regulation in endothelial cells

We next investigated cross-layer relationships to uncover underlying regulatory mechanisms. We used COJO-GCTA to perform conditional analysis and to identify independent association signals at each locus, followed by fine-mapping to construct 99% credible sets (**Methods**). We systematically assessed colocalization across molecular QTL layers, defining shared credible sets as those with a posterior probability ≥ 0.75 (**Methods**). This approach enabled us to link genetic effects on chromatin activity with effects on gene expression. In total, we identified regulatory features (**2,629** caQTL and **2,565** hQTL) for **2,064** genes and connected **4,638** out of **8,476** credible sets (55%) to at least one gene (**Fig. 2A; Table S6**). These acted either proximally by altering expression, or isoform usage of nearby genes (**Fig. 2B**), or distally via long range interactions bypassing the closest genes (**Fig. 2C**). Notably, **931** credible sets were associated with gene expression and/or isoform changes in more than one gene, highlighting both local and context-dependent regulatory complexity (**Fig. 2D-E**). The largest overlap was between hQTL and caQTLs (**2,681** colocalizing credible sets, 50% of all hQTLs and 45% of all caQTLs), while all other categories could be directly linked to at least a gene via eQTLs, isoQTLs, and/or sjQTLs. For eQTLs, **2,548** out of **5,646** were tested, for **1,248** we found colocalisation with caQTL and/or hQTL (**Fig. 2A**). To explore potential regulatory mechanisms mediating the effect of the shared hQTLs/caQTLs credible set, we performed transcription factor motif (TFBM) enrichment analysis. This identified enrichments for transcription factors central to endothelial biology and stress response, including AP-1 (JUN, FOS, FOSL1, FOSL2, JUNB) (Ma et al. 2024) (Wang et al. 1999), and ETS family members (ETS1, FLI1, FEV, ERG) (Meadows, Myers, and Krieg 2011) (Randi et al. 2009) (**Suppl. Table S7**). GO analysis of genes near h/caQTLs highlighted terms related to cell-to-cell signalling and adhesion (**Table S8**), consistent with these being poised enhancers under basal conditions. Furthermore, we observed that hQTLs and caQTLs not associated with eQTLs were located further away from transcription start sites (Welch Two Sample t-test; *p*-value < 2.2×10^−16^), supporting a role for distal enhancer regulation, in line with recent findings (**Suppl. Fig. S2**) (Arthur et al. 2025).

**Fig. 2.**
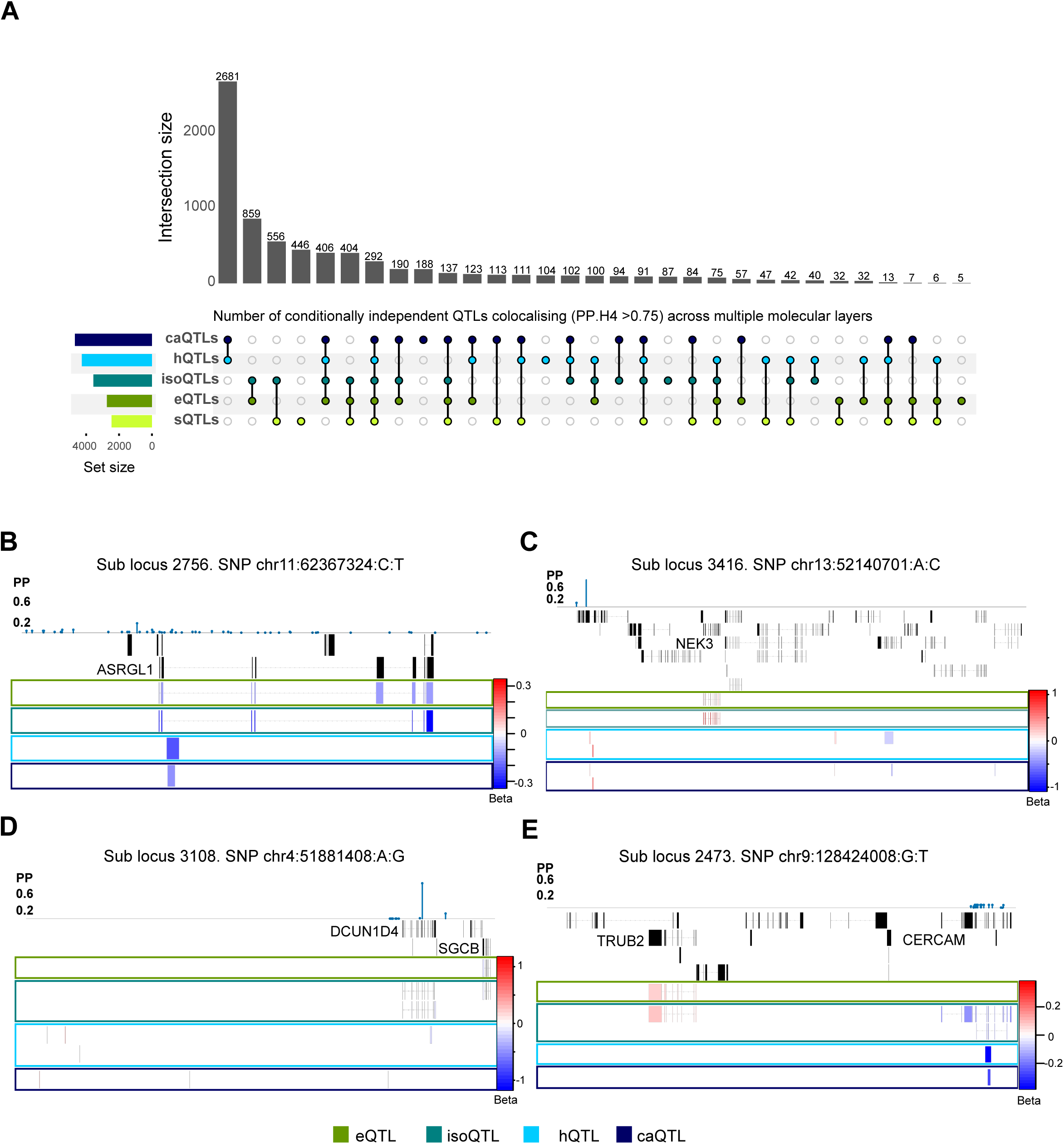
Shared Genetic Regulation of Gene Expression, Splicing, and Chromatin Activity. **A**. UpSet plot showing the number of credible sets shared between molecular QTLs. **B**. **Example of proximal credible set** (2756) with an effect on *ASRGL1* expression levels, transcriptional isoform usage, H3K27ac peak strength and open chromatin peak strength. **C**. **Example of distal credible set** (3416) with an effect on *NEK3* expression levels, transcriptional isoform usage, H3K27ac peak strength and open chromatin peak strength without affecting nearby genes. **D**. **Example of proximal credible set** (3108) with an effect on gene expression, *SGCB*, transcriptional isoform usage, *DCUN1D4* and *SGCB*, H3K27ac peak strength and open chromatin peak strength. **E**. **Example of credible set** (2473) acting both proximally, *CERCAM* transcriptional isoform usage, and distally, *TRUB2* expression levels and transcriptional isoform usage, with also effects on H3K27ac peak strength and open chromatin peak strength, without affecting other nearby genes.

### Pleiotropic cardiometabolic disease loci implicate vascular processes across multiple disease endpoints

Genome-wide association studies (GWAS) have identified thousands of common variants contributing to cardiometabolic disease risk. However, the cell types and mechanisms through which these variants act remain less well understood (Claussnitzer et al. 2020). We used the molecular QTL dataset to investigate the genetic architecture of diseases with established links to endothelial dysfunction. We looked for evidence of statistical overlap between the molecular QTLs and the GWAS summary statistics across a total of **2,961** credible sets associated with the following diseases: coronary artery disease (CAD; **204** loci), cardiovascular disease (CVD; **224** loci), type 2 diabetes (T2D; **203** loci), atrial fibrillation (AF; **119** loci), pulse pressure (PP; **620** loci), systolic blood pressure (SBP; **712** loci), diastolic blood pressure (DBP; **685** loci), haemorrhoidal disease (HD; **97** loci), pulmonary heart disease (PHD; **11** loci), venous thromboembolism (VTE; **13** loci), pulmonary arterial hypertension (PHA; **5** loci), phlebitis and thrombophlebitis (PTP; **9** loci), stroke (**24** loci) including the subtypes cardioembolic stroke (CES; **7** loci), large artery stroke (LAS, **2** loci) and ischemic stroke (IS; **26** loci), **Table S6 & S9**.

We first quantified pleiotropy across disease endpoints, thus defining their shared allelic architecture. Of **2,961** disease credible sets, **1,087** (37%) were shared between at least two diseases (**Fig. 3A**). Hypertension traits had the largest share (**n=931** colocalising loci between PP, SBP and DBP), consistent with their physiology. Additional statistical coincidence was observed between CVD and CAD (n=**53 loci**), T2D and CVD (n=**28 loci**) and VTE and PTP (n=**10 loci**), corresponding to pleiotropy rates of 12%-25%. To explore the shared biology, we assigned genes to disease loci using GREAT (McLean et al. 2010), and performed enrichment analysis. CAD-CVD shared loci were enriched for terms associated with blood vessel development (*p*=6.80×10^−6^), whereas PHD–VTE–PTP shared loci were enriched for blood coagulation (*p*=3.34×10^−11^) and wound healing (*p*=7.69×10^−9^; **Fig. 3B**).

**Fig. 3.**
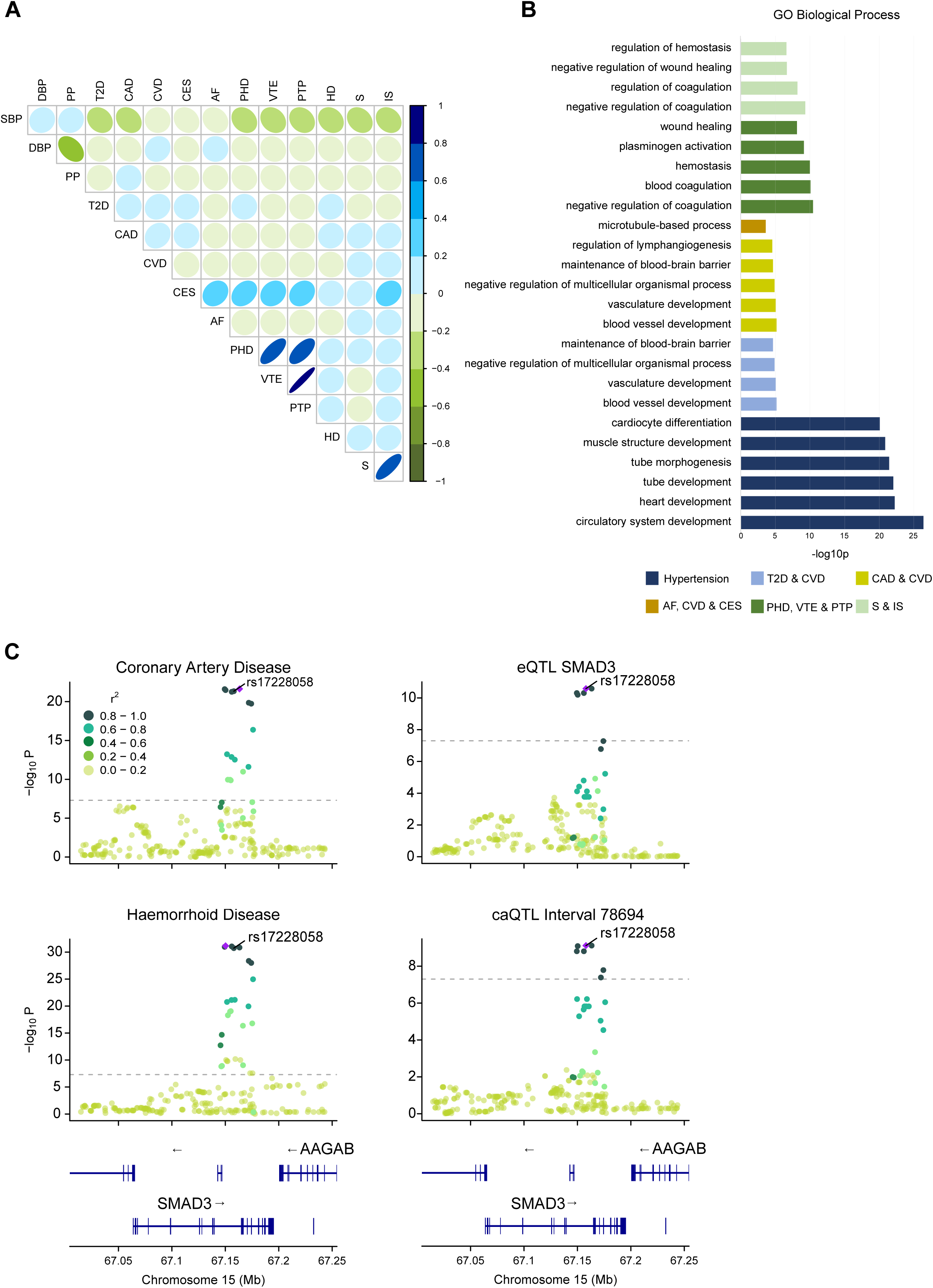
Cross-Trait Colocalisation of GWAS Signals Highlights Common Biological Pathways. **A.** Dot plot displaying the Pearson correlation calculated from the number of overlapping credible sets between disease GWAS. **B. Functional enrichment at pleiotropic disease loci.** Barplot showing -log10 *p*-values for GO terms obtained using genes shared disease loci **C. SMAD3 locus example.** Locus zoom plot displaying -log10 *p*-values for CAD, HD, SMAD3 eQTL and caQTL at the locus. LD information is calculated from rs17228058.

We next assessed the contribution of endothelial cells to disease outcomes by testing colocalization between GWAS loci and HUVEC molecular QTLs. In total, **815** disease credible sets colocalized with at least one QTL (**Suppl. Fig. S3A-C)**. The disease loci explained varied, ranging from 40% for PP (252/620 loci) to 14.3% for CES (1/7 loci), with only PAH and LAS having no colocalizing molecular QTL. Among complex traits with >100 loci, endothelial molecular QTLs signals explained 39% of SBP (277/712) and DBP (270/685) loci, 35% of CVD (79/224), 34% of T2D (70/203), 38% of CAD (77/202), and 29% of AF (34/119; **Table S6**). In total, this analysis linked **1,036** genes to at least one disease locus (**Suppl. Fig. S3D**). Contribution-wise caQTLs explained the largest number of loci, followed by isoQTLs (**Suppl. Fig. S3B**).

The *SMAD3* locus, previously implicated in CAD (Liu et al. 2018), illustrates the value of multilayered QTL integration. We confirmed colocalization of the CAD signal with HD (PPH4=0.99), as well as, with an eQTL (PPH4=0.99, beta=-0.19) and an isoQTL for *SMAD3* (ENST00000327367, PPH4=0.99, beta=-0.19). Fine mapping indicated an intronic caQTL(chr15:67149749-67150999; PPH4=0.99, beta=-0.17) (**Fig. 3C**). This contains binding sites for FOS, JUN and GATA2 (Hammal et al. 2022), and the risk allele rs17293632 (C>T; caQTL, *p*=8.11×10^−10^, decreased SMAD3 gene expression, decreased CAD risk, and increased HD risk) in perfect LD (r^2^=1) with the lead disease variant, predicted to alter JUN:FOS affinity (Kumar, Ambrosini, and Bucher 2017) (**Suppl. Fig. S3E**). These results provide mechanistic evidence linking a disease-associated locus to specific regulatory features in endothelial cells.

### Genetic risk for cardiovascular disease intersects endothelial and immune cell transcriptional programs

Chronic inflammation is a contributing factor for cardiovascular disease (Willerson and Ridker 2004) and targeting inflammation reduces severe cardiovascular events (Ridker et al. 2017). However, the understanding of the molecular mechanisms underlying these observations remain incomplete. Monocytes, neutrophils, and T lymphocytes interact directly with the vascular endothelium during atherogenesis, plaque destabilisation, and vascular repair (Phillipson and Kubes 2011) (Kwee et al. 2018) (Medrano-Bosch et al. 2023), yet the role of genetic variation in shaping these processes in a cell-type-specific manner has not been fully resolved.

To investigate the interplay between endothelial and immune cells, we compared endothelial eQTLs and hQTLs with corresponding data in three immune cell types, monocytes, neutrophils, and naive CD4 T lymphocytes (Chen et al. 2016). We performed colocalisation between eQTL and hQTL for three immune cell types and the same 16 disease GWAS. A comparable number of disease loci were explained across the 4 cell types for both eQTLs and hQTLs (**Fig. 4A; Table S10**).

**Fig. 4.**
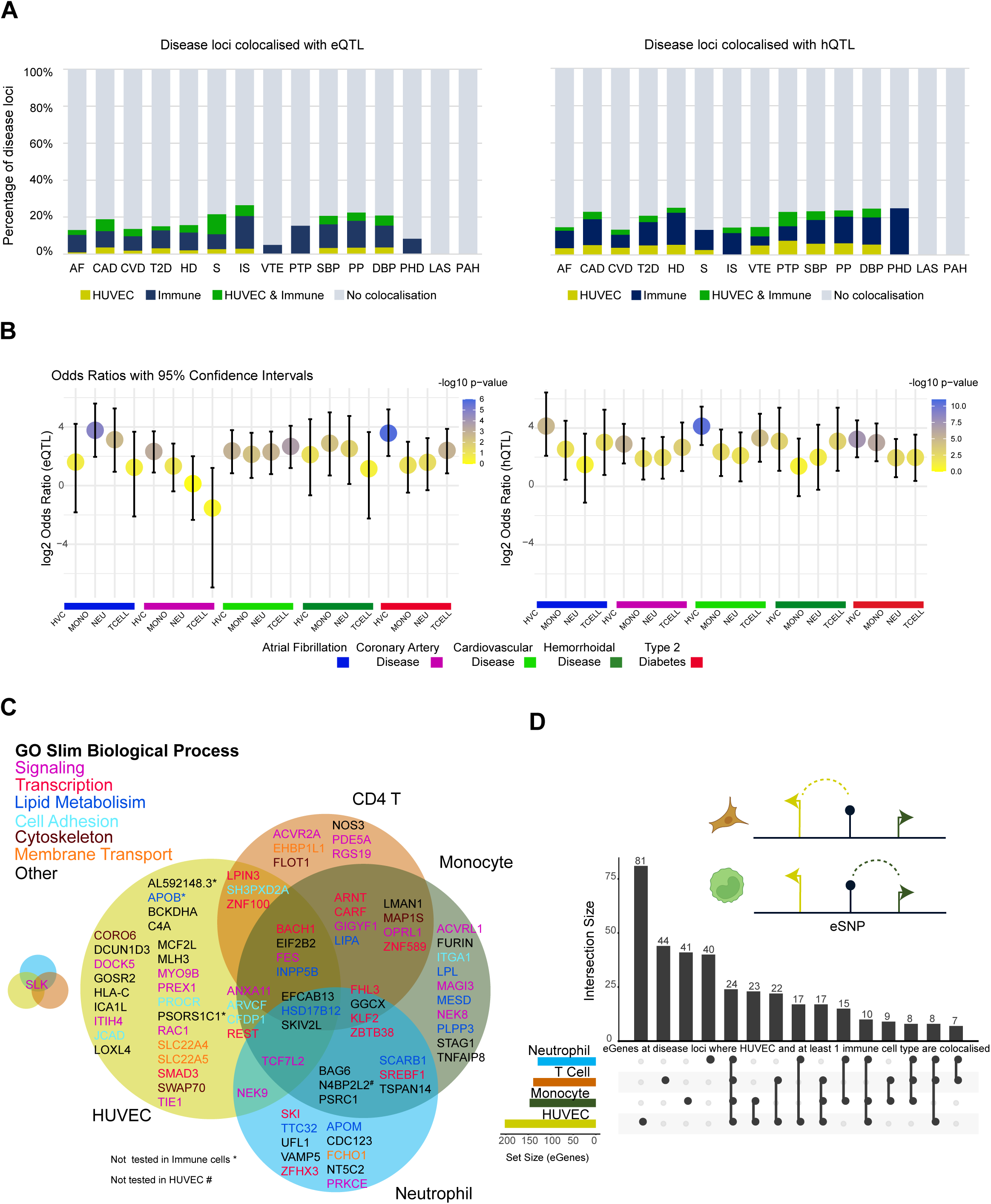
Colocalised GWAS Loci Reveal Convergent Immune and Endothelial Regulatory Mechanisms. **A**. Left; Stacked barplot displaying the percentage of disease loci colocalised with an eQTL with either HUVEC (darkyellow), at least 1 immune cell type (navy) or HUVEC shared with at least 1 immune cell type (green). Right; Stacked barplot displaying the percentage of disease loci colocalised with an hQTL with either HUVEC (darkyellow), at least 1 immune cell type (navy) or HUVEC and at least 1 immune cell type (green). **B**. Dot plot displaying the odds ratios of colocalised loci between GWAS and eQTLs (left) and hQTLs (right) from four cell types (HUVECs, monocytes, T cells, neutrophils), based on Fisher’s exact test. Each point represents the strength of enrichment (odds ratio) for a given cell type. The analysis tests the contribution of cell type specific colocalisations in explaining disease loci. P-values indicate the statistical significance of enrichment. Results are shown for five representative GWAS. **C.** Proportional venn diagram illustrating the overlap between candidate CAD genes and the cell types (HUVEC, monocytes, neutrophils, and CD4 T cells) in which we observed eQTL colocalisation with disease loci. For 42 loci, the colocalizing signal was found in HUVECs, 40 in monocytes, 26 in neutrophils, and 29 in CD4 T cells **D.** UpSet plot displaying the composition of eGENES at colocalised disease loci where each locus is explained by a HUVEC eQTL and at least 1 immune cell type.

We combined eQTLs and hQTLs credible sets for each cell type and quantified the overlap between HUVEC and immune cells (neutrophils, monocytes, CD4 T lymphocytes) (**Table S11**). GREAT analysis showed that credible sets unique to immune cells were enriched for adipose tissue and heart morphology-related terms, credible sets unique to endothelial cells were enriched for vascular phenotypes and endocrine function terms, while shared sets showed no biological enrichment (**Suppl. Table S12**). Across all four cell types, the fraction of disease loci explained ranged from 18% (n=371; SBP) to 32% (n=328; PP). The largest percentage was contributed by immune cell QTLs, followed by shared signals, and then endothelial-specific signals (**Table S12**). The latter ranging from 20 (23%) of the explained CAD loci to 5 (11.5%) of the AF loci.

We identified 1,501 disease eQTL colocalising pairs across the four cell types (HUVEC n=534, Monocyte n=671, Neutrophil n= 637 and CD4 T cell n=584). We also identified colocalizations for n=789 HUVEC, n=653 monocytes, n=551 neutrophils and n=482 CD4 T cells hQTLs with at least one disease locus. The majority of these hQTLs were cell-type specific, with up to 35% unique to HUVECs. Cell-type specific hQTLs tended to map farther from the nearest transcription start sites compared to shared hQTLs (Welch Two Sample t-test *p*=3.43×10^−7^, **Figure S4**). Cell type specific disease colocalisations, and their contribution to the number of loci explained differed across cell types, the strongest eQTL associations were between monocytes and AF, HUVEC and CAD, HUVEC and T2D, and monocytes and HD. For hQTLs the strongest association was observed between HUVEC and CVD (**Fig. 4B** and **Table S6 & S10**). We intersected the 1,501 gene/disease GWAS loci pairs with the 420 CAD-associated genes curated by Schnitzler *et al*. (Schnitzler et al. 2024) and found an overlap of 86 (**Fig. 4C**). Additionally, Schnitzler *et al*. also included 41 high-confidence prioritised genes (variant-to-gene framework), of these we detected 18: 7 in HUVEC (*CFDP1, GOSR2, PREX1, SH3PXD2A, SLK, SMAD3, SWAP70*), 5 in monocytes (*CFDP1, FURIN, N4BP2L2, PLPP3, TSPAN14*), 3 in neutrophils (*N4BP2L2, SLK, TSPAN14*), and 3 in CD4 T cells (*NOS3, SH3PXD2A, SLK*). Several of these are well-established CAD genes with strong functional plausibility; *NOS3* encodes endothelial nitric oxide synthase, central to vascular tone and endothelial health; *PLPP3* regulates vascular integrity and shear-stress responses; *FURIN* processes proprotein substrates, including PCSK9, thus implicating lipid metabolism; and *SMAD3* is a key mediator of TGF-β signaling, implicated in vascular remodeling and fibrosis. Finally, for curated genes not prioritised in this framework, we observed several instances where the potential effector cell was one of the three immune cell types. Interesting examples include: *TNFAIP8, SCARB1, LPL, ZFHX3, ITGA1, PRKCE, APOM, SKI, FCHO1,* and *BAG6*. The presence of these well-established causal genes across multiple cell types supports a model in which both endothelial and immune mechanisms contribute to CAD risk, with some loci acting through both compartments and others in a cell-type-specific manner.

Due to extensive co-regulation in the genome, eQTLs often associate with more than one gene (eGENE) (Tambets et al. 2024), We therefore assessed the concordance between eGENES at cell-shared eQTL loci. Overall, 59% of HUVEC eGENES were also observed in at least one immune cell type (**Fig. 4D**), and only 11–22% of eGENES were unique to each cell type (e.g. 40/366 in neutrophils, 81/366 in HUVECs; **Fig. 4D**). We next examined allelic effects, and whether shared eQTLs had the same directionality of effect across cell types. While most shared eQTLs had concordant directionality, a minority displayed opposite effects between endothelial and immune cells, or even between immune cell types, e.g. the *NEK9* locus (**Suppl. Fig. S5A-D**).

The locus containing *ZNF664* and *CCDC92* illustrates the complexity of cross-tissue regulation. It has been associated with CAD, immune, and metabolic phenotypes (Dastani et al. 2012) (Kraja et al. 2014) (W. Zhao et al. 2017) (**Suppl. Fig. S6A-B**). Here, rs7975482 is an eQTL for *ZNF664* and *CCDC92* in HUVEC and naive CD4 T cells, but only for *ZNF644* in neutrophils (**Suppl. Fig. S6C-D**). hQTLs identified regulatory regions in CD4 T cells (chr12:123999065-124009132) and monocytes (chr12:123989419-123991461), but only the CD4 region colocalized with the eQTL. Within this enhancer, rs12311848 (LD r^2^=1 with rs7975482), was previously targeted by CRISPRa in teloHAEC cells by Wünnemann et al. (Wünnemann et al. 2023), leading to CCDC92, but not ZNF664, upregulation.

A second example is the *IL6R* locus. Interleukin 6 (IL-6) is a pro-inflammatory cytokine central to immune activation (Tanaka, Narazaki, and Kishimoto 2014), and independent variants at this locus have been associated with autoimmune and cardiovascular diseases (Zhang et al. 2022). eQTL analysis in whole blood (Ferreira et al. 2013) (Tokolyi et al. 2025), and purified immune cell types (Chen et al. 2016), determined that rs2228145 alternative allele result in a change in splice junction usage and the production of the soluble form of the receptor. We observed colocalization between multiple HUVEC molecular QTLs (including the known sjQTL), and CAD (PPH4=0.84) and AF (PPH4=0.85), and between a caQTL with CAD (PPH4=0.97) and AF (PPH4=0.95). Immune cells percent spliced in (psiQTL) confirmed consistent effects across these and disease traits (**Suppl. Fig. S7 & S8**), with neutrophil hQTLs (1:154399555:154447432) colocalizing with CAD (PPH4=0.941) and AF (PPH4=0.942). These examples underscore the importance of examining gene expression effects in the context of chromatin annotation information.

### Genetic mapping of endothelial barrier ECIS traits identifies genes underlying functional phenotypes variation

Genetic association frameworks can be applied to cell function phenotypes. We performed a GWAS using Electric Cell-substrate Impedance Sensing (ECIS) measurements of impedance, resistance, and capacitance in response to wounding and to thrombin stimulation. Functional Principal Component Analysis was applied to each assay-measurement pair, and the resulting scores were used for GWAS analysis (**Methods**). Associations across wounding and thrombin assays were highly correlated (**Suppl. Fig. S9A**). No genome wide significant associations (*p-val*≤1×10^−8^) were found for the thrombin assay (**Suppl. Fig. S10**). One locus, rs113309218, reached genome-wide significance for wounding resistance and impedance (**Suppl. Fig. S9B, S10**), with clear genotype associated response profiles (**Suppl. Fig. S9B**). This variant lies within the second intron *CDH9* (**Suppl. Fig. S9B**), which encodes a type II classical cadherin, mediating calcium-dependent cell-cell adhesion. Although *CDH9* is predominantly expressed in the brain and not in steady-state HUVECs, this locus may influence endothelial barrier responses through context dependent effects. Given the modest sample size, we next explored variants associated at nominal significance (*p*<5×10^−5^; LD-pruned at r^2^ ≥0.8, n=267; **Table S13**). GREAT analysis of these loci revealed enrichment for cell-adhesion pathways and mouse gene knockout phenotypes related to myocardial function (**Fig. 5A**).

**Fig. 5.**
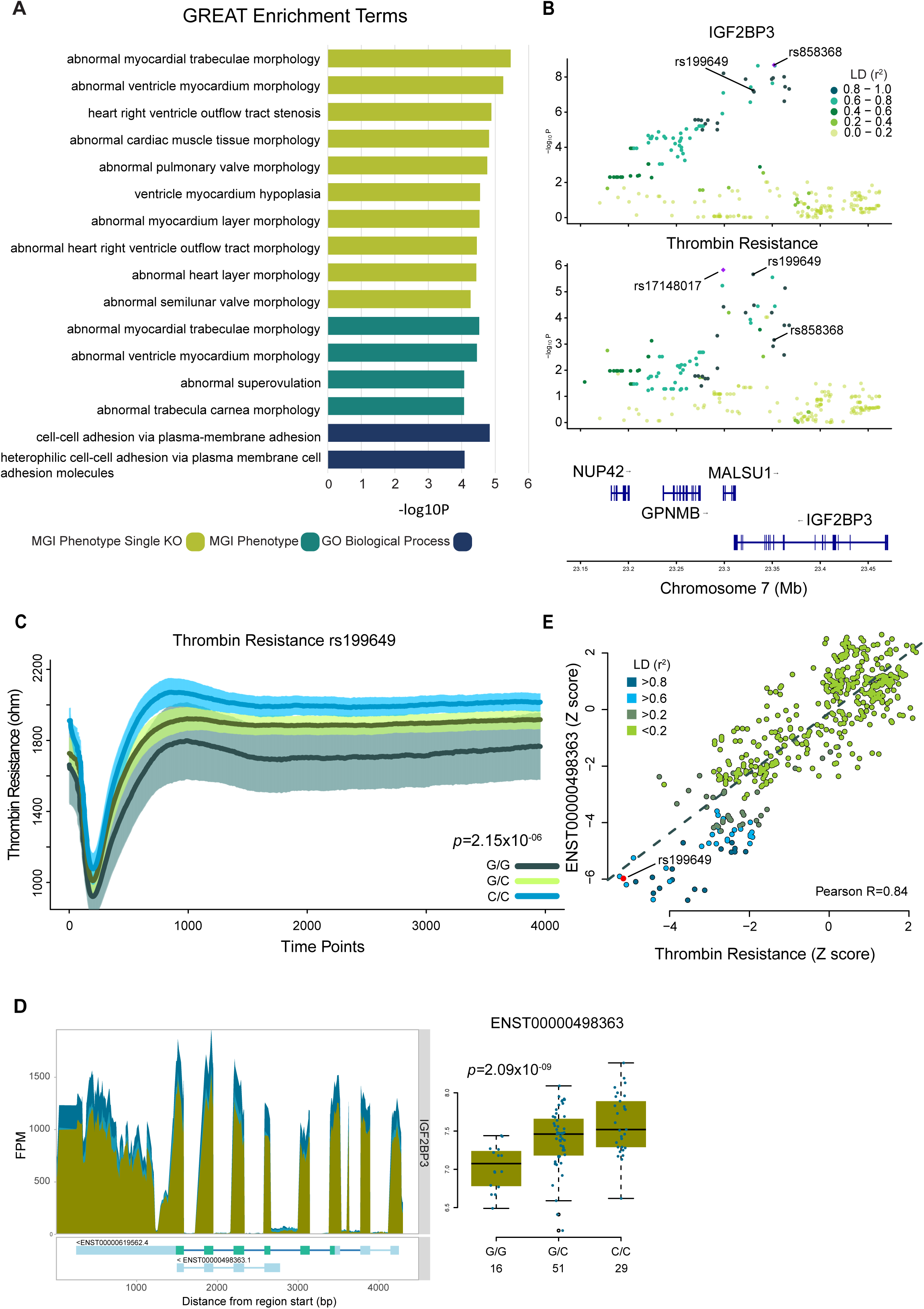
Integrated Genetic Analysis Identifies IGF2BP3 as a Regulator of Endothelial Barrier Function. **A**. Barplot displaying enriched functional categories using GREAT tool. ECIS SNPs for all tests were pruned LD r^2^>0.8 and used as input. **B.** Locuszoom plots for; isoQTL transcript ENST00000498363 at the *IGF2BP3* gene in HUVEC (top) and thrombin resistance (bottom). The top shared associated SNPs for each trait are highlighted. **C**. ECIS data for thrombin resistance in ohms (y-axis) and time (x-axis) coloured by genotype status for rs199649. Thick dark lines are the mean thrombin resistance values for each genotype, shaded areas are the standard error with 95% confidence intervals. **D.** Left; Transcript coverage plot at the *IGF2BP3* gene with signals from representative donors; REF (dark sky blue), HET (blue) and ALT (dark olive green). Right; Gene expression boxplots for the short isoform ENST00000498363, split by genotype status for SNP rs858368. **E**. Scatter Plot displaying the SNP-trait Z scores between ENST00000498363 and Thrombin Resistance. LD information is calculated from the lead eQTL rs858368.

Eight ECIS-associated loci overlapped with molecular QTLs ((r²>0.8; **Table S14**). Amongst these, we found an isoQTL (rs858368, *p*=2.09×10^−9^, beta=0.28) for *IGF2BP3* (Insulin-like growth factor 2 mRNA-binding protein 3) in high LD (r^2^=0.84) with rs199649, associated with thrombin resistance (*p*=2.15×10^−6^; **Fig. 5B-E**).

We intersected ECIS loci with disease GWAS and identified two AF loci in LD (r^2^>0.8): rs8032930 at the *CTXND1* locus (wounding impedance *p=*4.52×10^−5^; AF *p=*5.80×10^−7^) and rs3003311 at the *ADSS2* locus (thrombin impedance *p=*2.17×10^−6^ and AF, *p=*6.01×10^−5^; **Suppl. Fig. S11**). *CTXND1* is an eQTL in atrial appendage and heart left ventricle (GTEx Consortium 2013)(Leblanc et al. 2024), and *ADSS2* is a molQTL in our dataset (**Suppl. Fig. S11**). Although underpowered, this analysis highlights candidate loci and genes involved in the genetic regulation of endothelial barrier function. Integration with GWAS and molQTL datasets suggests that endothelial barrier phenotyping may help resolve pleiotropic GWAS signals in future, larger studies.

### HUVEC molecular QTLs inform on target prioritisation, drug repurposing, and type of therapeutic modulation

Therapeutics raised against targets supported by genetic evidence have an increased likelihood of successful approval (Finan et al. 2017). We analysed our dataset to identify targets supported by genetic evidence. We selected credible sets in the eQTL and isoQTL datasets colocalizing with disease GWAS loci, as these correlated the directionality of the effect on disease with expression levels. We identified 357 genes (including lncRNAs; **Suppl. Table S15**) that were filtered to identify those with the desirable characteristics of a druggable target, i.e. encoding for an enzymatic activity or a cell surface protein, having a crystal structure, having a high confidence ligand, being a known target of drug/antibody/small molecule (Finan et al. 2017). Of these, 71 had at least one of these properties, including 23 genes linked to known compounds. Ten genes are targets, or have demonstrated interactions, with 17 approved therapeutics, while 13 others are the targets of compounds either in clinical trials or discontinued. In all but one gene/compound combinations (*PDE3A*/pentoxifylline), the indication for which the compound is used, or being developed for, is unrelated from the disease GWAS in which the colocalisation was found, thus highlighting new genetically supported drug repurposing opportunities. The remaining 42 genes, filtered to retain only those with a single significant eGene colocalized with the disease locus, represent candidates for therapeutic prioritization. For these, the sign of the correlation between the directionality of the molecular QTL and disease-associated allele informs whether agonist or antagonist modulation would be appropriate (**Fig. 6**).

**Fig 6.**
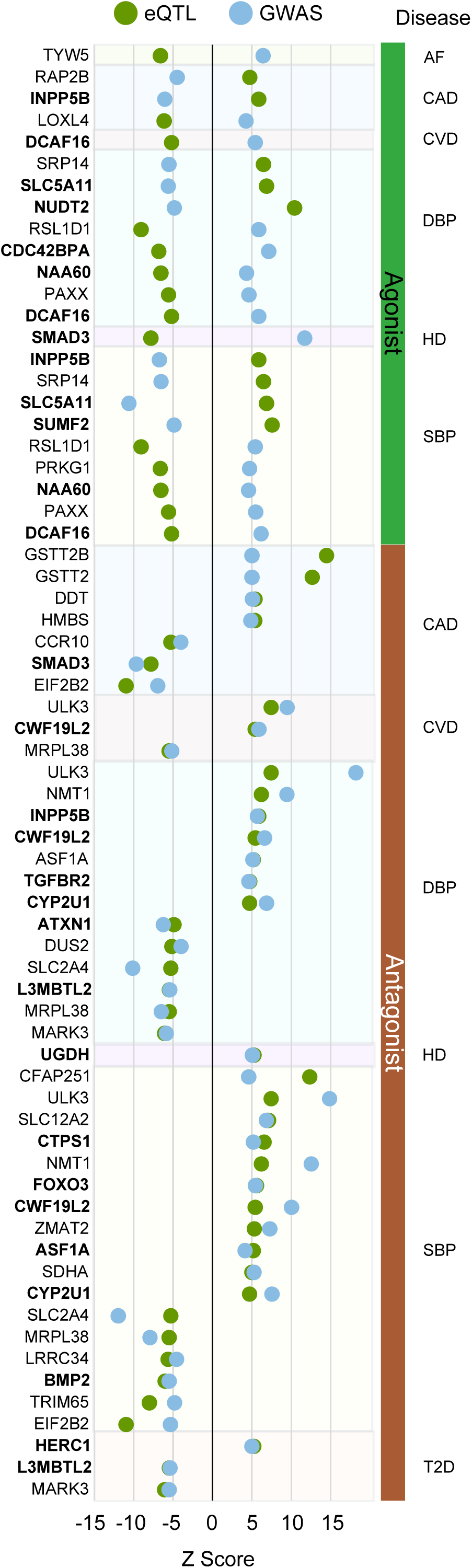
HUVEC eQTLs Inform on Directionality of Therapeutic Modulation. Dot plot showing a list of potential target genes that have potential as therapeutic agents. Genes are segregated based on whether the gene expression correlates with disease risk (antagonist) or is anti-correlated to disease risk (agonist). Z-scores are calculated from the top eQTL SNP. Gene names highlighted in bold are those where there is only 1 significant eGene colocalized with disease at that locus.

## Discussion

We generated a new dataset by profiling both molecular (RNA-seq, H3K27ac ChIP-seq, ATAC-seq), and functional assays in HUVEC samples derived from 98 unselected cord blood donors. The dataset was used to determine the contribution of common genetic variation to endothelial cell traits, deriving information on genetic variants influencing gene expression, splicing, active chromatin, chromatin occupancy and functional phenotypes in endothelial cells. These were then used to identify the genes, and molecular mechanisms, underlying GWAS loci for cardiometabolic diseases having endothelial dysfunction as a common feature.

Altogether, we identified regulatory features for 8,214 genes (5,646 genes regulated by eQTLs, 5,295 by isoQTLs and 2,687 by sjQTLs). We also profiled cis-regulatory elements using H3K27ac ChIP-seq and chromatin accessibility, together identifying 8,541 regulatory regions under genetic control. By applying statistical colocalisation we were able to nominate single credible sets as having an effect on two, or more, data layers. The largest group of colocalizing credible sets was that of hQTLs and caQTLs, likely representing either poised regulatory elements used in response to stimuli (Fairfax et al. 2014), or for which we lacked power to detect expression changes for the linked genes.

Interestingly, for 931 credible sets we found effects (eQTL/isoQTL) on more than one gene, likely because of alterations of the local chromatin topology and/or in changes in the ability to access the transcription and splicing machinery for the involved genes. This is relevant when using this approach to identify the genes behind the association at disease GWAS loci. To this end, we used this data to explain disease GWAS loci acting through endothelial cells, and to connect them to genes. We explained **815** disease GWAS loci, representing between 14% and 40% of all known loci for each of the 16 diseases analysed, and between 29% and 40% for studies with more than 100 identified loci. This is a considerable increase over the average 23% of disease GWAS loci explained by eQTLs (Mostafavi et al. 2023) and it is explained by a combination of the additional molecular QTLs (isoQTLs, sjQTLs, hQTLs, caQTLs), and by the use of a disease relevant cell type. Not surprisingly, given the regulatory nature of GWAS loci, caQTLs and hQTLs had the largest contribution. However, the contribution of isoQTLs and sjQTLs, which allowed us to quantify the genetic effects on alternative splicing, was also substantial.

We further quantified the extent to which genetic effects were pleiotropic across different diseases. While the traits linked to hypertension (PP, SBP, DBP) had the largest amount of loci overlap (>80%), for other disease pairs we observed pleiotropy ranging from 12% to 25%. Pleiotropic loci shared between disease pairs had specific GO term enrichments (**Table S12**), highlighting common developmental pathways, but also the prominent role of the coagulation cascade in diseases with endothelial dysfunction as common denominator.

The inclusion of molQTL data from immune cells allowed us to identify shared and cell type specific genetic regulatory mechanisms linking vascular and immune pathways to cardiovascular disease. The number of explained disease GWAS loci was similar across cell types, we found that the immune cells molecular QTLs explained the largest number of GWAS disease credible sets, while HUVEC specific eQTL contributed significantly to the number of CAD and T2D loci explained. eGENES were more likely to be shared across cell types, whereas hQTLs had higher tissue specificity, in agreement with the concept that enhancers have a higher degree of tissue specificity than gene expression. Shared eGENES had functions linked to signalling, lipid metabolism, cell adhesion, cytoskeleton, membrane transport, and transcription.

Lastly, we developed a framework to identify cell function QTLs. While statistically underpowered, our analysis identified 267 nominal associations, whose credibility is supported by the enrichment for relevant GO terms and high linkage disequilibrium with known disease loci. We identified *IGF2BP3* (**Fig. 5B**), a member of the insulin-like growth factor family, known to play a role in mediating cardiovascular risk and are key proteins in endothelial function and homeostasis. It has also been shown that its expression exerts protective effects by alleviating endothelial dysfunction caused by ox-LDL (Gao et al. 2024). This approach should be further explored to identify loci acting as phenotype modifiers, and to gain further understanding of the underlying genes. Altogether, our results provide a quantification of non-communicable diseases heritability acting through endothelial cells. We have identified pleiotropy between diseases and within different cell types in mediating disease susceptibility, adding a further layer of complexity to the understanding of these diseases’ aetiology.

### Limitations of the study

Although our study provides new insights on the role of endothelial cells in non-communicable diseases, the following limitations should be considered. While the study allowed for the detection of common variation effects, a larger sample size is required to capture the effects of rare genetic variants. This would also allow us to draw robust conclusions on genetic variants altering cell phenotypes. The study also lacked samples harvested after different stimuli, so we could only speculate upon the role of enhancers under genetic control, but not linked to a gene. Furthermore, the complete landscape of the effects of genetic variation cannot be obtained without considering endothelial cells from other vascular beds, as well as, without considering other ancestries.

## Materials and Methods

### Data availability

All scripts used are publicly available at https://github.com/maxozo/HUVEC. Raw data are publicly available, under managed access in accordance with the ethical approval, at: RNA-seq (EGAS00001006476), ATAC-seq (EGAS00001006606) and ChIP-seq (EGAS00001006208).

A detailed description of all methods is available in Supplementary Information.

## Acknowledgments

This work was supported by British Heart Foundation (BHF) grants PG/13/25/30014, PG/14/69/31032, PG/16/63/32307, PG/09/015/26991, FS12/8/29377, PG/20/10270, PG/20/10387, RE/18/1/34212, RG/13/14/30314, RG/16/13/32609, RG/19/9/34655, FS/SBSRF/20/31005 and FS/18/53/33863. Wellcome Trust (206194), Rosetrees Trust (CM374), the BHF Cambridge Centre for Research Excellence and the NIHR Cambridge Biomedical Research Centre (NIHR203312). This research was funded in whole, or in part, by the Wellcome Trust 220540/Z/20/A. For the purpose of Open Access, the author has applied a CC BY public copyright licence to any Author Accepted Manuscript version arising from this submission. SY is supported by the National Medical Research Council (MOH-001229 and MOH-001479) of Singapore, the National University of Singapore/National University Health System (NUHSRO/2022/004/Startup/01). DGM is supported by the Leicester British Heart Foundation Research Excellence Award (RE/24/130031) and the van Geest Foundation Heart and Cardiovascular Diseases Research Fund, and was awarded a BHF Accelerator Early Careers Researcher Interdisciplinary Fellowship and pump-priming funding from the Leicester British Heart Foundation Accelerator Award (AA/18/3/34220). This study was supported by the National Institute for Health and Care Research Exeter Biomedical Research Centre. The views expressed are those of the author(s) and not necessarily those of the NIHR or the Department of Health and Social Care. We acknowledge the contributions of May Hu and Feri Torabi for technical assistance. We thank members of Emma Davenport’s laboratory for fruitful discussions and feedback.

---

**Supplementary Figure 1.**
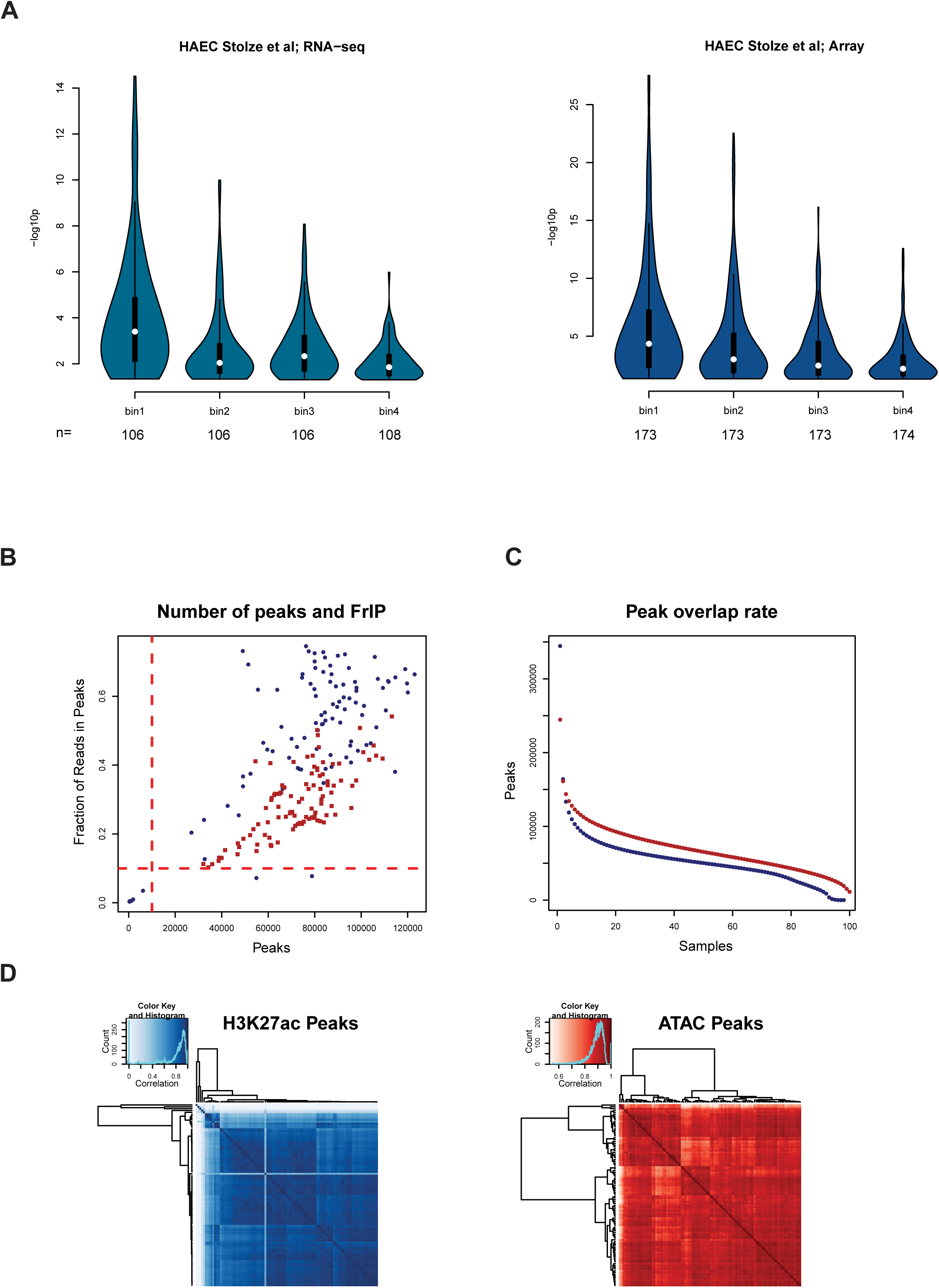
Validation of HUVEC eQTLs in an external dataset. ChIP-seq and ATAC-seq quality control. **A.** HUVEC eQTLs were ranked and sorted into 4 bins. Violin plots display the corresponding -log10p values for LD matched eQTLs from Stolze et al. **B. Quality control fraction reads in peak**. Scatterplot with the number of peaks identified in a sample on the x-axis and the fraction of mapped reads within those peaks on the y -axis. H3K27ac ChIP-seq in blue and ATAC-seq in red. **C. Quality control number of shared and unique peaks.** Overlap rate plot displaying the sum of peaks identified across all donors decaying to the number of peaks shared within all individuals. H3K27ac in blue and ATAC in red. **D Quality control, sample correlation.** Heatmap showing the peak overlaps correlation for H3K27ac (blue) and ATAC (red).

**Supplementary Figure 2.**
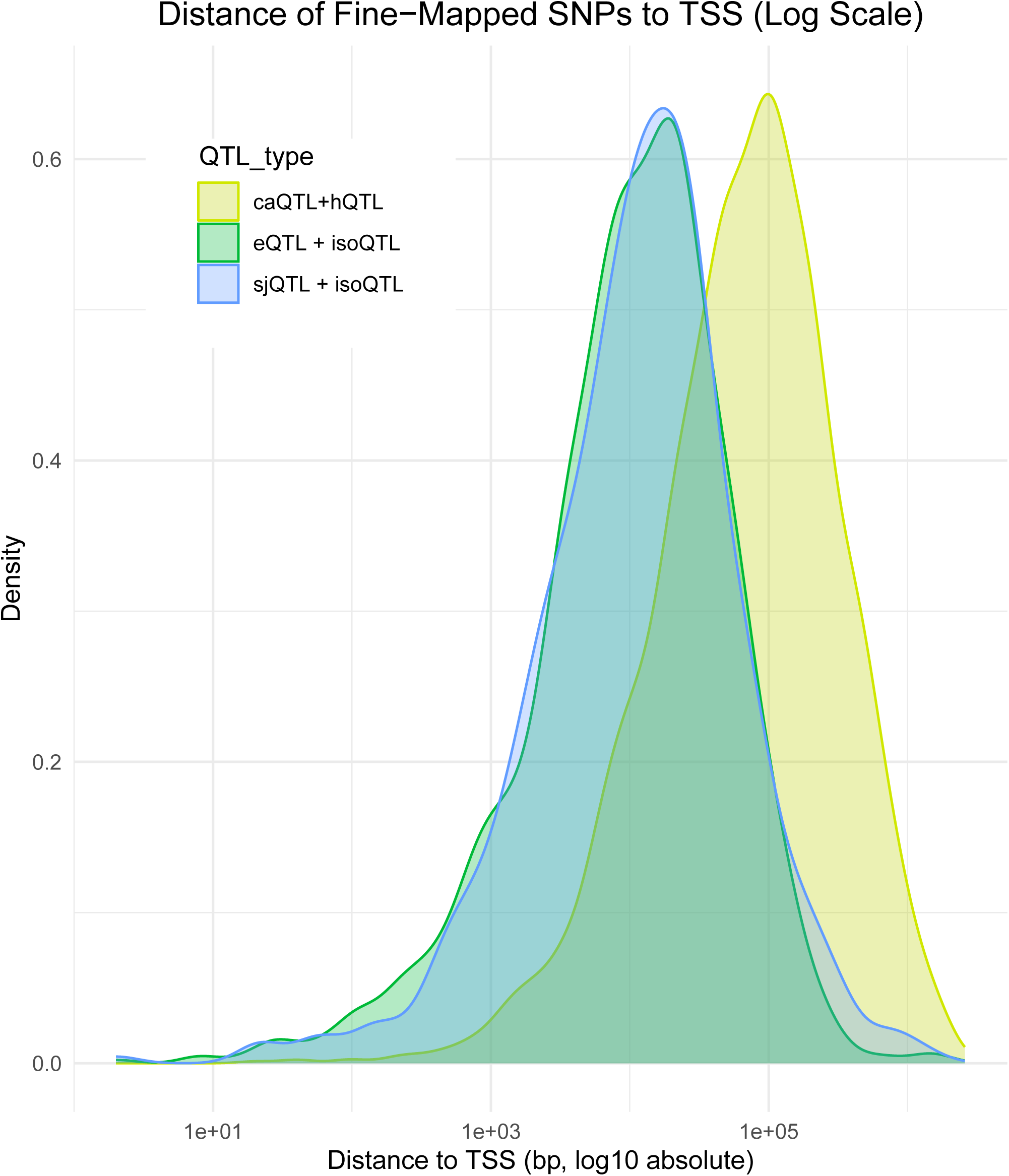
Fine Mapped Chromatin QTLs Colocalise with Variants Located Distally to Transcription Start Sites. Density plot showing genomic distance of the lead fine mapped variant in relation to the transcription start site of the nearest protein coding gene. Three categories of lead variant are shown; caQTL that colocalise with hQTL and without evidence of colocalisation with sjQTL, isoQTL or eQTL (yellow), sjQTL that colocalise with isoQTL without evidence of colocalisation with chromatin QTLs (blue) and eQTL that colocalise with isoQTL without evidence of colocalisation with chromatin QTLs (green). Chromatin QTLs not associated with transcriptional QTLs tend to be located farther from TSSs (Welch Two Sample t-test *p*-value < 2.2×10^−16^ for both eQTL and sjQTL comparison to chromatin QTL), consistent with regulation at distal enhancer elements.

**Supplementary Figure 3.**
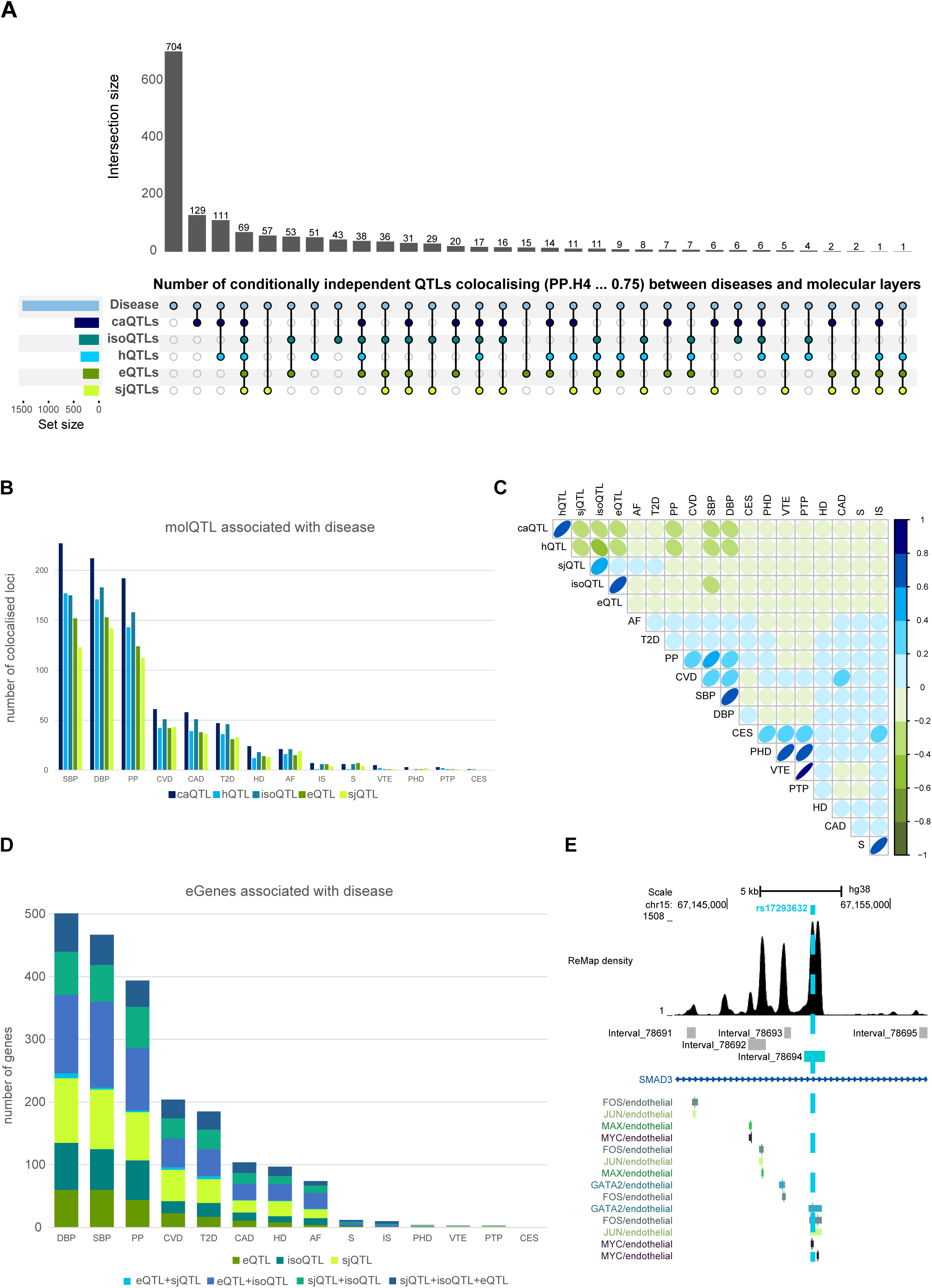
Colocalised Endothelial QTL Signals Across Transcriptomic and Epigenomic Layers. **A**. UpSet plot displaying molQTLs and disease associated credible sets. **B**. Barplot displaying number of colocalized credible sets by molQTL type and each disease GWAS. **C**. Pearson correlation calculated from the number of overlapping colocalised credible sets by trait and molQTL. **D.** Stacked barplot displaying the number of eGENES at each colocalised credible set by trait and by the molQTL data type. **E.** Genome browser view with ChIP-seq track data from ReMap and displaying the location of the caQTL (light blue) within the *SMAD3* intron and the position of the variant rs17293632 (light blue dashed line).

**Supplementary Figure 4.**
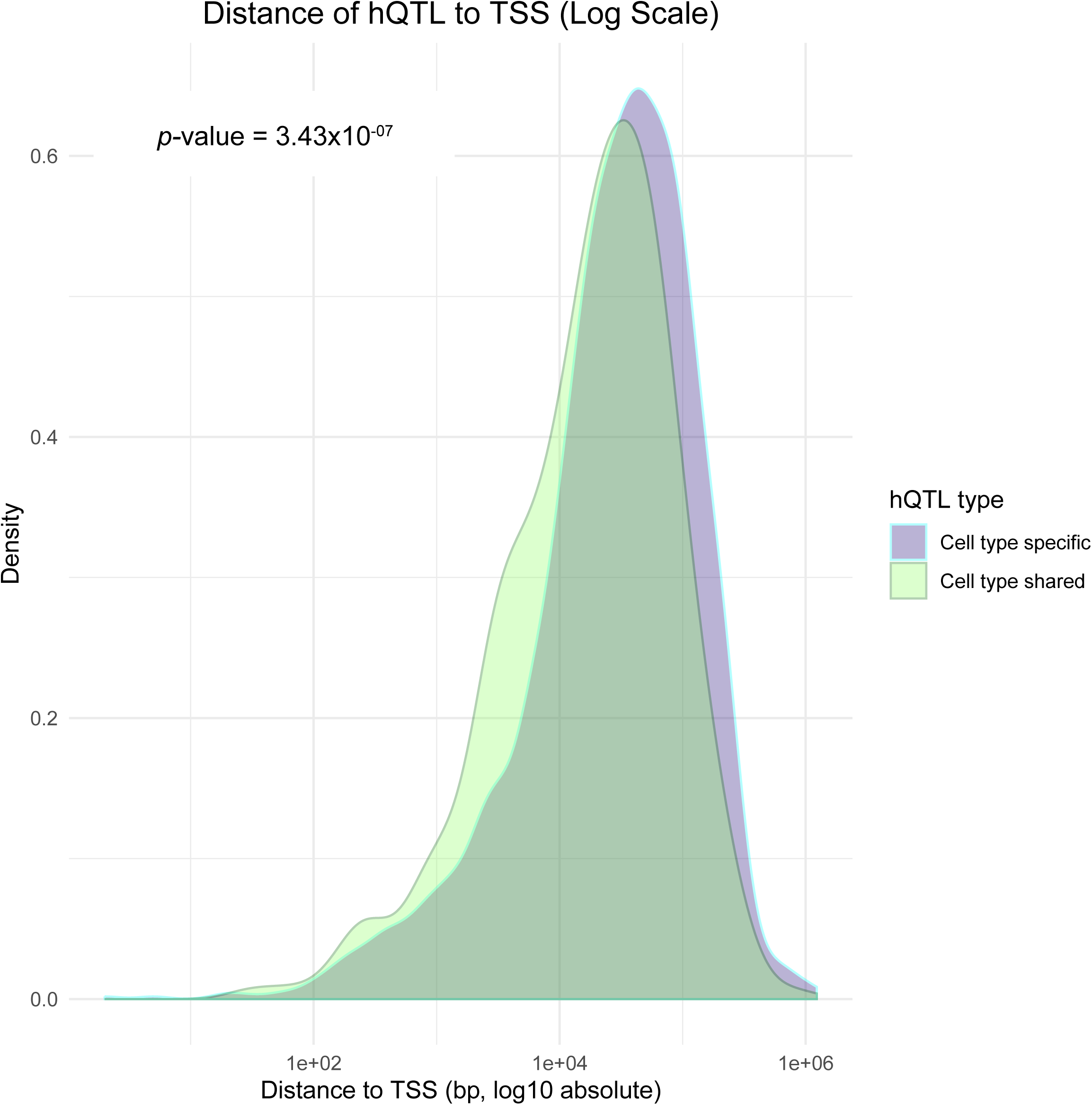
Cell Type Shared Associated hQTL Regions are Closer to Transcription Start Sits than Cell Type Specific. Density plot showing genomic distance of disease colocalized hQTL’s in relation to the transcription start site of the nearest protein coding gene. Two categories are shown; hQTL regions that are cell type specific (blue) and hQTL regions that overlap in at least 2 cell types (green). A merged hQTL peakset for both studies was created (1bp overlap) and the distance was calculated from the centre of the hQTL to the nearest TSS. *p*-value calculation was performed using a Welch Two Sample t-test.

**Supplementary Figure 5.**
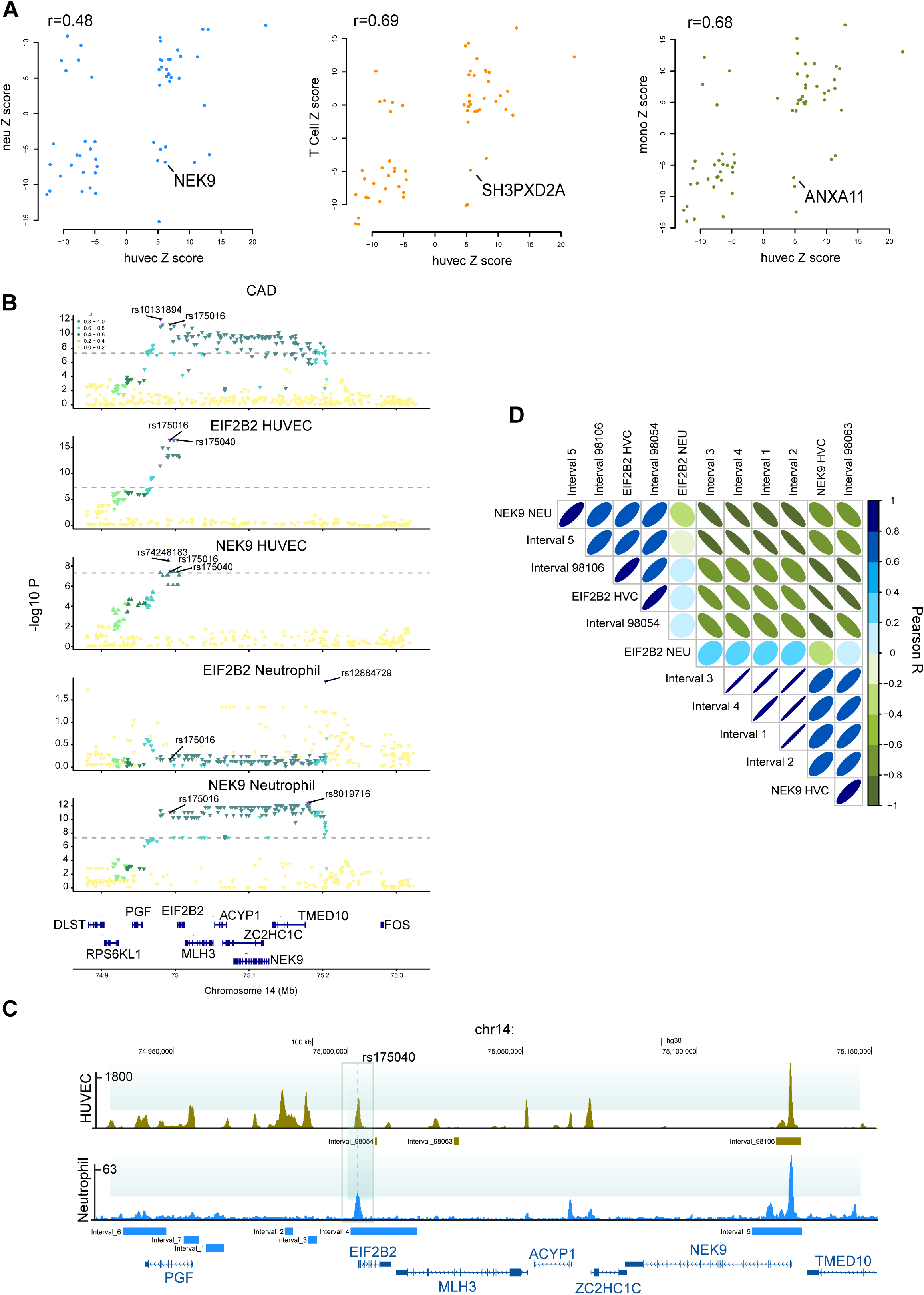
Limited Divergence in eQTL Effects Between Endothelial and Immune Cells. **A.** Scatter plots displaying Z scores for the allelic effect on gene expression. The y-axis shows results for neutrophils (dodger blue), CD4 T cells (orange), and CD14+ monocytes (olive green), plotted against HUVECs on the x-axis. We identified 14 genes in neutrophils, 10 in CD4 T cells, and 9 in monocytes where the alternative allele had an opposite effect on gene expression compared to HUVECs. Pearson correlation ( R ) is shown in the upper left. **B.** LocusZoom plot of the genomic region surrounding the *EIF2B2* and *NEK2* genes. Summary statistics are shown for coronary artery disease (top), and eQTLs for *EIF2B2* and *NEK9* in HUVECs and neutrophils as indicated above each plot. Triangle direction indicates the alternative allele’s effect on the respective phenotype. Linkage disequilibrium (r²) values are based on rs175016 from the 1000 Genomes Project. **C.** Genome browser view of H3K27ac coverage in HUVECs (dark yellow) and neutrophils (light blue). Genomic intervals used in hQTL analyses from each study are displayed below each track. The fine-mapped lead variant rs175040 is marked with a dashed line. **D.** Pearson correlation plot showing correlations of Z scores between molQTLs across cell types at the *NEK9* locus.

**Supplementary Figure 6.**
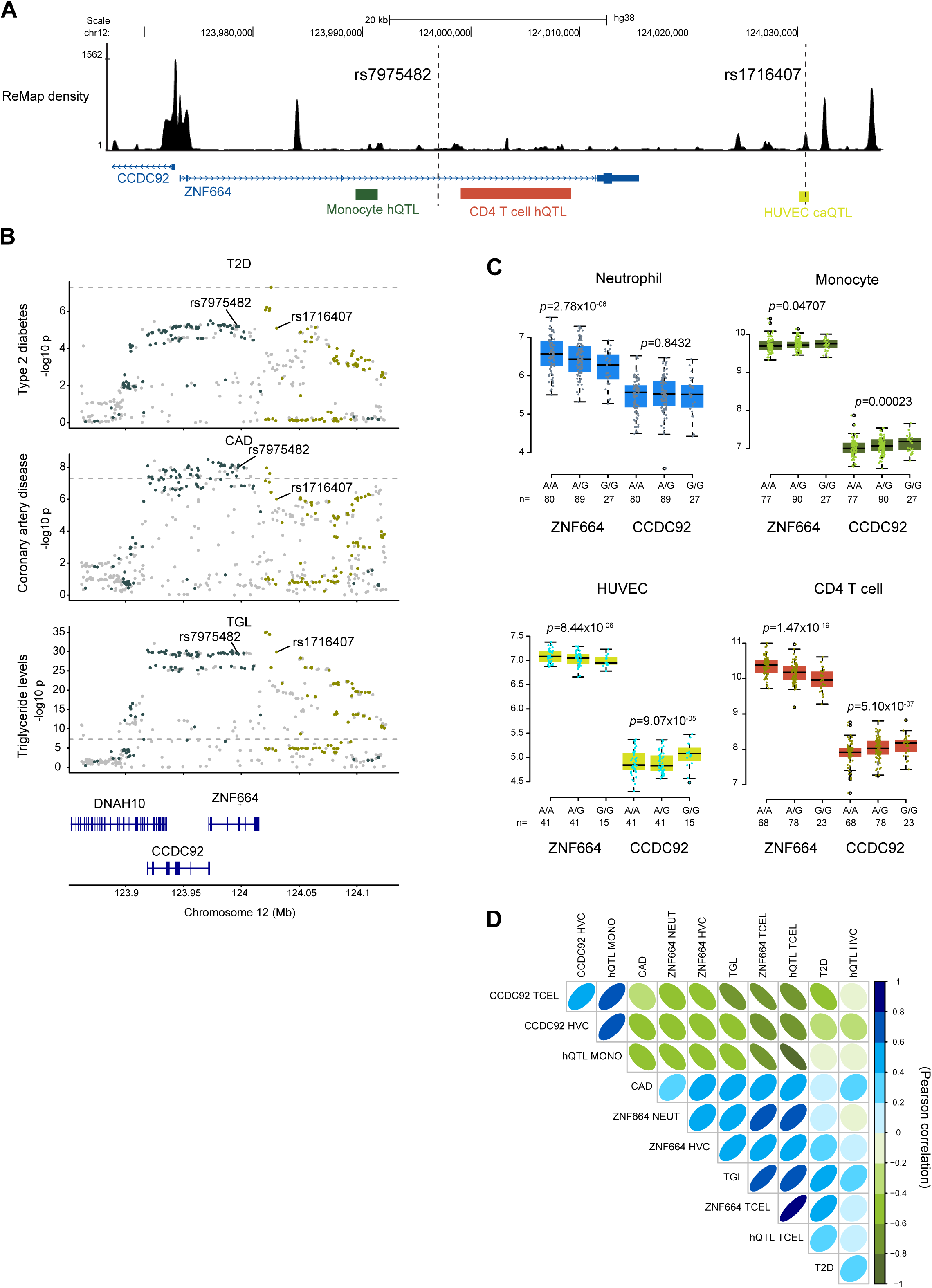
Multiple Regulatory Variants at the ZNF664 Locus Exhibit Distinct Cell-Type Activity. **A**. Genome browser region at the *ZNF664* gene with track data for Endothelial ChIP-seq data from ReMAP and the locations of monocyte hQTL, CD4 T cell hQTL and HUVEC caQTL beneath. Dashed line highlighting the position of lead eQTL SNP rs7975482 and lead caQTL SNP rs1716407. **B**. Locus zoom plots displaying windows around the *ZNF664* and *CCDC92* genes. Top; Type 2 diabetes, middle, Coronary artery disease and bottom Triglycerides. Variants in LD r^2^ >0.2 with rs7975482 are marked in dark slate gray. Variants in LD r^2^ >0.2 with rs1716407 are marked in dark yellow. **C.** Boxplots displaying the gene expression levels for *ZNF664* (left) and *CCDC92* (right) for HUVEC (lime green), CD4 T cell (dark orange), neutrophil (blue) and monocyte (olive green). Donors are segregated by genotype status for the variant rs7975482. **D**. Pearson correlation plot between Z scores for each molQTL and the three selected GWAS traits from B.

**Supplementary Figure 7.**
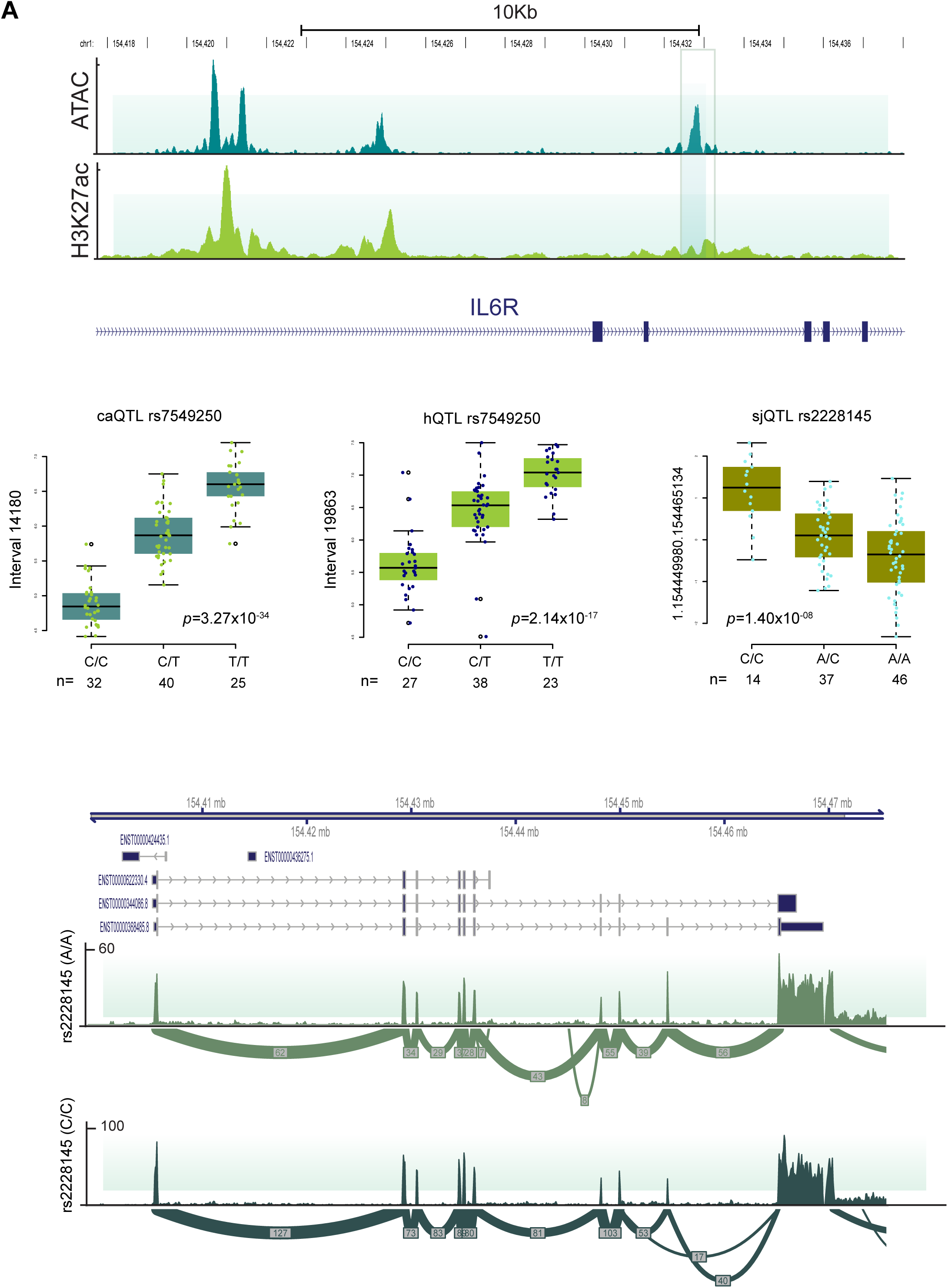
Chromatin and Splice Junction Genetic Effects at the IL6R Gene. **A**. Illustrative genome browser view highlighting the location of the enhancer QTL between the 3rd and 4th exons of the *IL6R* gene, (shaded area). **B**. Quantitative trait boxplots split by genotype status for caQTL (deepskyblue) hQTL (yellowgreen) and sjQTL (olive). **C.** Sashimi plots split by genotype status for the *IL6R* gene. Top is representative of splice junction usage in carriers of the reference allele for variant rs2228145. Bottom is representative of splice junction usage in donors carrying the alternative allele.

**Supplementary Figure 8.**
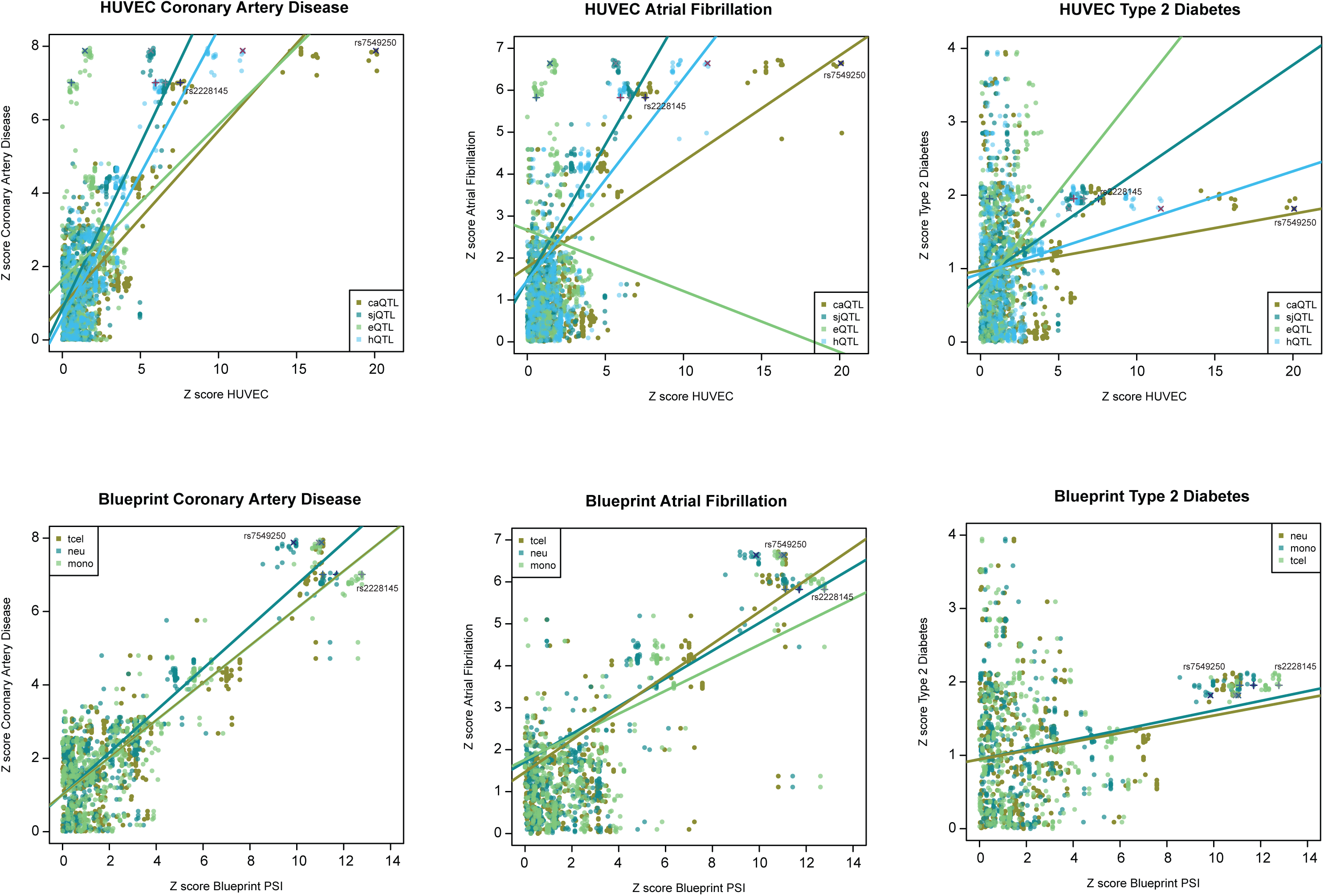
Shared Splicing Effects at the IL6R Gene among Endothelial and Immune Cells Associates with Coronary Artery Disease and Atrial Fibrillation. Scatterplot of Z-scores for variants at the *IL6R* locus between disease (y-axis) and molecular QTL (x-axis). Top row is HUVEC molecular QTL data; caQTL(olive), sjQTL(aqua), eQTL (seagreen) and hQTL (lightskyblue). rs2228145 is marked on the plots by (+) and rs7549250 by (x). Type 2 diabetes is representative as a negative association. Bottom plots are as above, where QTL data is percent spliced in (psi) from three immune cell types; CD4 naïve T cell (seagreen), Neutrophil (olive) and CD14 monocyte (aqua).

**Supplementary Figure 9.**
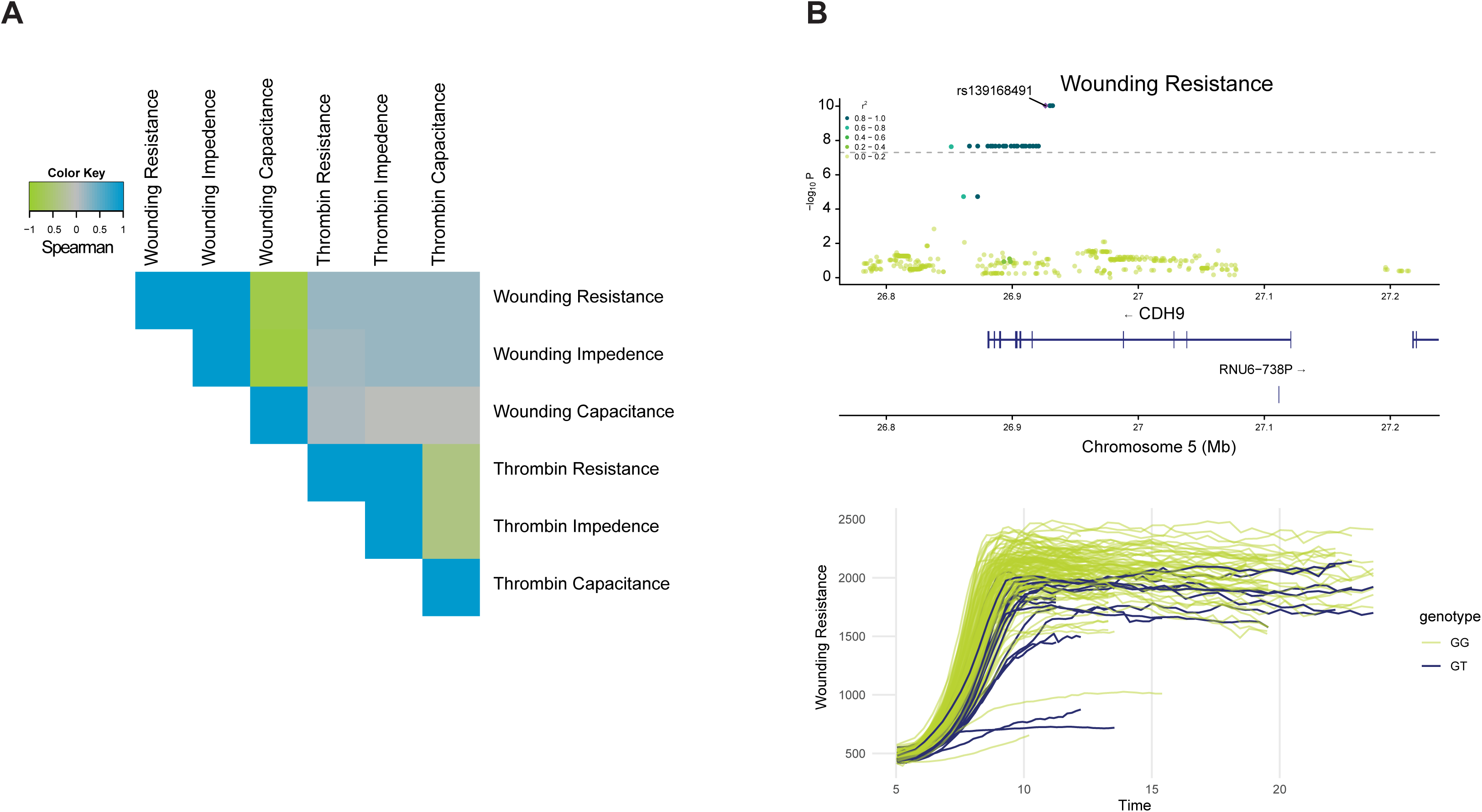
Endothelial Barrier GWAS Identifies Physiologically Relevant Loci. **A.** Spearman correlations between Z-score for all nominally significant SNPs (p<5×10−5) in the 6 GWAS tests. **B.** Locus zoom plot for the genome wide significant hit at the *CDH9* locus in Wounding Resistance test (top). ECIS electrical measurements for the 92 donors coloured by genotype (bottom). Olive green for reference genotype (GG) and navy the heterozygote (GT). There were no alternative allele (TT) carriers in our test cohort.

**Supplementary Figure 10.**
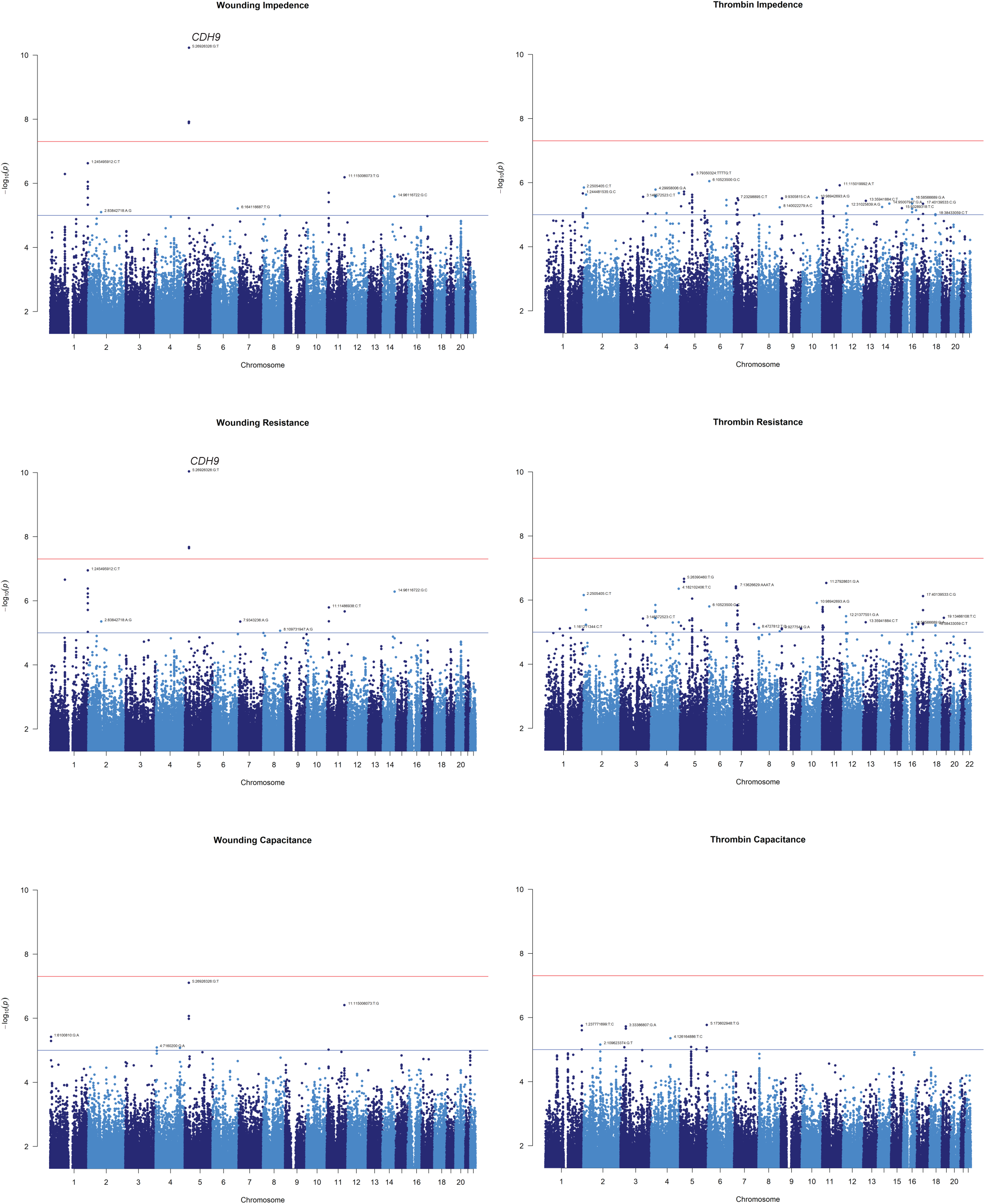
GWAS of Endothelial Functional Traits. **A**. GWAS Manhattan plots for the 6 GWAS performed across 2 traits; wounding and thrombin exposure and 3 measurements; capacitance, impedance and resistance.

**Supplementary Figure 11.**
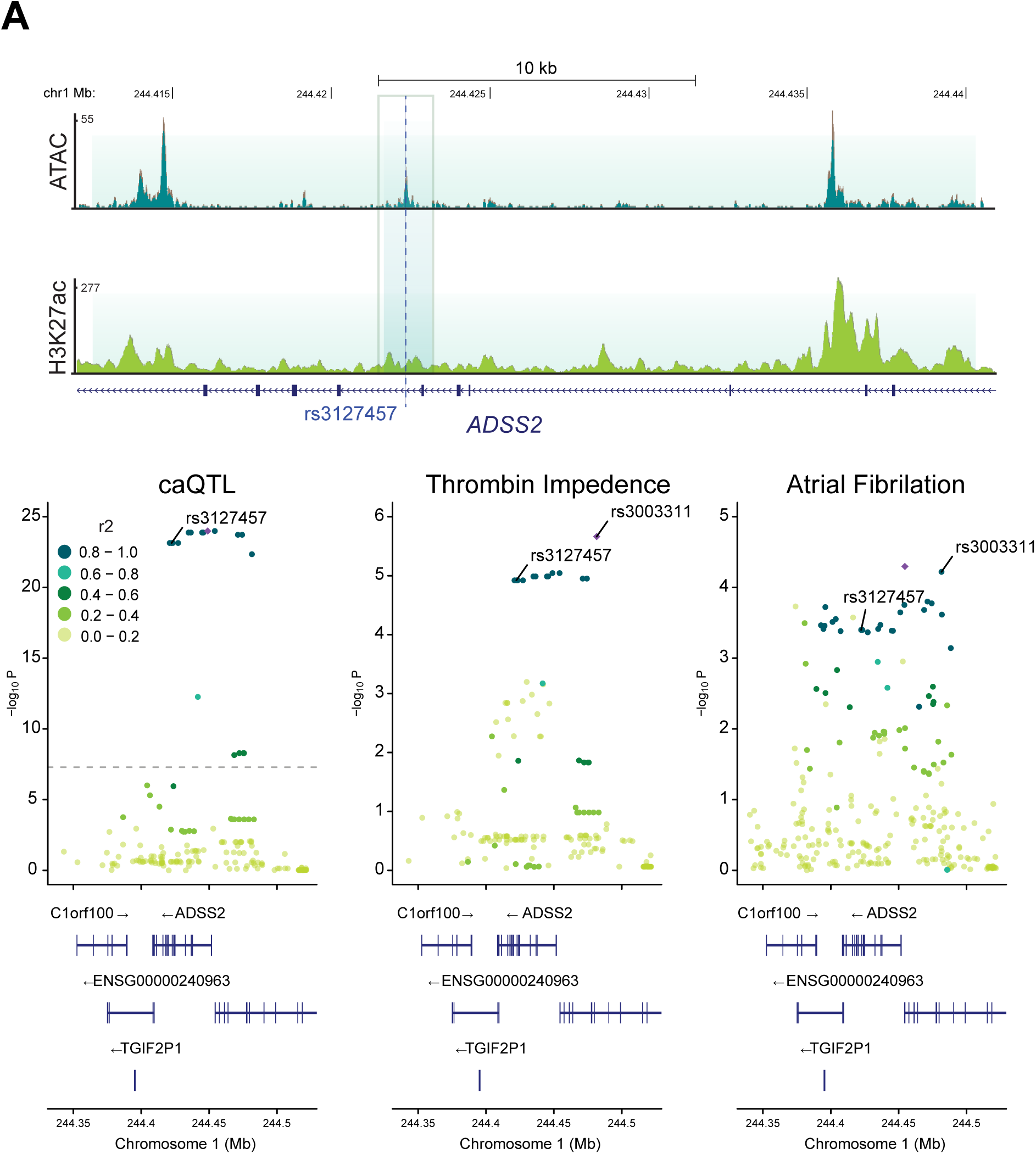
Endothelial Barrier GWAS Overlaps with Putative Disease Relevant Loci. **A.** Genome browser view with representative ATAC and H3K27ac tracks at the *ADSS2* locus (top) with location of variant predicted to disrupt transcription factor binding highlighted by dashed line and caQTL region highlighted by shaded area. Locus zoom plots for caQTL, Thrombin Impedance and GWAS for Atrial Fibrillation (bottom).

## Materials and Methods

### Sample collection

Human umbilical vascular endothelial cells (HUVEC) were isolated from umbilical cords provided by the Anthony Nolan foundation (https://www.anthonynolan.org/) after informed consent of the parents. Each cord was flushed with PBS and then treated with collagenase as previously described (Baudin et al. 2007) to harvest HUVEC. Cells were seeded on 0.2% gelatine-coated T75 flasks in M199 (Sigma M4530) supplemented with 15% FCS, 1% pen/strep, 4.5ug/ml ECGS (Sigma E2759), 10U/ml heparin (Sigma H3393), 2.5ug/ml FGF-acidic (Peprotech, 100-17A) and 2.5ug/ml thymidine (Sigma 89270), split 1:2 and then frozen once confluent. One hundred samples, of Northern European ancestry, were thawed and used in all subsequent experiments. Suppl. Table S1 contains the metrics of the samples in this study. Cells were maintained in Endothelial Cell Growth Medium 2 (EGM2, Promocell, C-22111) supplemented with 10% Foetal Bovine Serum (FBS) in 5% CO2 at 37 C and used at passage 2.

### Genotype data

DNA was extracted from each cord and genotyped using the Infinium Global Screening Array-24 2.0 BeadChips (Illumina) as already described (Solomon et al. 2022). Imputation was performed using the TOPMED imputation server, and the 1000 Genomes phase 3 mixed population (Das et al. 2016) served as the reference panel. To filter variants we used PLINK (v1.9) (Purcell et al. 2007) using the following parameters –geno 0.05 –hwe 1e-06 –maf 0.01 –mind 0.1, after which a total of 8,671,174 variants remained. PLINK was also used to calculate the principal components of the filtered data and to recode the genotype data in 0,1,2 or change the format of the data to VCF where required.

### RNA-Sequencing library preparation

Total RNA was extracted using Qiagen RNeasy kit, including optional on column DNAse digestion, as per manufacturer’s instructions. RNA integrity values (RIN) were obtained using Agilent Bioanalyzer RNA Nano 6000 kit, a median RIN value of 9.7 was obtained for all samples, however one sample had a value of 4.2 and was included in the study, and one further sample was removed. Samples were quantified with QuantiFluor RNA System from Promega UK Ltd on a BMG FLUOstar Omega plate reader. Ribosomal RNA was depleted using the Kapa HMR rRNA depletion kit and libraries made using the NEB Ultra II RNA custom kit (NEB #E7775) on an Agilent Bravo WS automation system. PCR set-up using KapaHiFi Hot start mix and Eurofins dual indexed tag barcodes. Post PCR plate purified using Agencourt AMPure XP SPRI beads on Caliper Zephyr liquid handling platform. Libraries were quantified with Biotium Accuclear Ultra high sensitivity dsDNA Quantitative kit and BMG FLUOstar Omega plate reader. Samples were pooled at equimolar concentration and diluted to 2.8nM before sequencing on an Illumina NovaSeq 6000 S4 paired end 100 bp without XP.

### ATAC-Sequencing library preparation

Nuclei were prepared from ∼100,000 HUVEC cells with ice cold lysis buffer (10 mM Tris-HCl, pH 7.4, 10 mM NaCl, 3 mM MgCl2, 0.1% v/v IGEPAL CA-630) on ice for 15 minutes. Nuclei were resuspended in transposase buffer (100 mM Tris-HCl, pH 8.00, 50 mM Magnesium Chloride) and tagmented with Illumina Nextera Kit (FC-121-1031) at 37oC for 30 minutes. Index sequences were applied using the Illumina dual index kit (FC-131-2001) with 12 cycles of PCR.

Samples were pooled and diluted to 2.8nM before run on an Illumina NovaSeq 6000 S2 paired end 50 bp without XP.

### Chromatin Immunoprecipitation library preparation

Cells were fixed with 1% formaldehyde at a concentration of approximately 1 million cells/ml. Fixed cell preparations were washed and stored re-suspended in PBS at 4°C prior to lysis and sonication. Sonication was performed in a Diagenode PicoRuptor for 25 cycles of 30 seconds on, 30 seconds off in a 4oC water cooler. Samples were checked for sonication efficiency using the criteria of 150-500bp, by Agilent DNA bioanalyzer. Immunoprecipitation was carried out as previously described (Schmidt et al. 2009) with minor modifications. Protein A Dynabeads (Invitrogen) were coupled with 5µg of H3K27ac antibody (Diagenode C15410196). Sonicated lysate (1-2 million cells) was then added to the bead/antibody mix and incubated at 4oC overnight. ChIP-DNA bound beads were washed for five repetitions in cold RIPA solution.

Elution and crosslink process reversal was carried out at 65oC for five hours. 2µl RNase was added to ChIP-DNA and incubated at 37oC for 30 minutes, followed by 2µl of Proteinase K treatment at 55 oC for 1 hour. 1:1.8 ratio of Ampure beads (Beckman Coulter, A63881) were added to the DNA followed by two cold 70% ethanol washes. Immunoprecipitated DNA was eluted in 50µl of elution buffer. Illumina sequencing libraries were prepared using NEBNext Ultra II reagents. Amplification was performed using Kapa HiFi master mix (Kapa Biosystems KK2602) for 16 cycles of PCR, followed by a 0.7:1 Ampure XP clean-up. In addition 2 background control libraries each prepared using whole cell extract pooled from 4 randomly selected donors; WCE1: 588, 591, 590, 563 and WCE2: 1044, 1092, 1095, 1233. Libraries were sequenced across 4 individual lanes of an Illumina NovaSeq 6000 100 bp paired end.

### Bioinformatic analyses

#### RNA-sequencing

We processed raw fastq files with the NF-core RNA-Seq pipeline (v3.6) (https://github.com/nf-core/rnaseq), using STAR (2.7.6a) for alignment and both RSEM (v1.3.1) and Salmon (v1.5.2) for quantification, adhering to default configurations. Our analysis included mapping to the GRCh38 release 99 human reference genome from Ensembl. This pipeline ensured rigorous quality control, integrating adapter trimming (Trim Galore!), read quality assessment (FastQC), and quantification (featureCounts), complemented by comprehensive reporting (MultiQC).

#### ChIP-sequencing

The raw fastq files generated were processed using the nf-core/ChIP-seq (https://github.com/nf-core/chipseq ) Nextflow pipeline using default settings in conjunction with Singularity (version 2.6.0). Samples were mapped to the human reference genome version GRCh38 release 99 obtained from Ensembl. To summarise, the pipeline utilises tools such as BWA for alignment, MACS2 for peak calling, deepTools for quality control and data visualisation, and featureCounts for quantification.

#### ATAC-sequencing

The raw fastq files generated were processed using the nfcore/atacseq (https://github.com/nf-core/atacseq) nextflow pipeline using default settings. Samples were mapped to the human reference genome version GRCh38 release 99 obtained from Ensembl. The nf-core/ATAC-seq pipeline uses similar processing steps as described for the nf-core/ChIP-seq pipeline in the previous section but with additional steps specific to ATAC–seq analysis, including removal of mitochondrial reads.

#### Molecular QTL analyses

We conducted eQTL analysis using the QTLight pipeline ( https://github.com/wtsi-hgi/QTLight ) across multiple omics datasets, including bulk RNA-seq, transcript RNA-seq, splicing, ChIP-seq, and ATAC-seq. In each analysis, genotype-phenotype mapping files were used, and a 1 Mb cis-window around the transcription start site (TSS) or peak region was applied to identify cis-regulatory variants. TensorQTL was employed for association testing, with 2 genotype PCs to account for population structure.

For RNA-seq, aggregated transcript-level counts were used as the phenotype, while for transcript RNA-seq, transcript-aligned reads from the nf-core RNAseq pipeline were used. In both cases, DESeq2 was applied for normalisation, and no filtering was applied for MAF or expression thresholds. The analyses followed the same process for cis-eQTL mapping, with a 1 Mb window around the TSS.

For splicing QTL analysis was performed using Leafcutter (de-novo assembler quantifying exon-intron excision, v0.2.9) to identify splicing events from bulk RNA-seq data. After generating junction files, splicing events were clustered, and intron usage counts were extracted. The raw intron usage counts were then normalised using a two-step process: row-wise scaling followed by quantile normalisation (qqnorm), which transformed the distribution to approximate a normal distribution. This normalised data was used as the phenotype for downstream association testing. A 1 Mb cis-window was applied around the cluster end location.

For both ChIP-seq and ATAC-seq QTL analyses, narrow peaks were called using the nf-core ChIP-seq and nf-core ATAC-seq pipelines, respectively, with peak calling performed using MACS2 (narrow peaks, gsize = 2.7e9). In both cases, reads were filtered to remove duplicates, blacklisted regions, and non-canonical reads. Peak counts were normalised using DESeq2, and peak sets were used as the phenotype for downstream association testing. A 1 Mb cis-window was applied around the end of the peak intervals, similar to the approach used for transcription start site (TSS) analysis, to detect variants affecting chromatin accessibility (for ATAC-seq) or H3Kac protein-DNA binding (for ChIP-seq).

### Electric Cell-substrate Impedance Sensing (ECIS) thrombin response and wounding assays

Cell proliferation, endothelial barrier permeability, and migration of HUVEC were measured in duplicate by trans-epithelial/endothelial electrical resistance (TEER) on an ECIS Theta machine. Each donor cell line was cultured to the same passage (P2) and seeded on two identical 8W10E ECIS arrays, which were measured in parallel. A HUVEC line obtained from Lonza (cc2519, lot No. 0000192485) was expanded and aliquots of the same passage seeded on each array as a positive control. On each array a well was left empty as a negative control and to measure the drift in baseline measurement.

Briefly, electrodes were first coated with fibronectin (Corning, 356008) followed by L-cysteine (Fisher Scientific) preparation. HUVECs were seeded at 20,000 cells per well in Endothelial Cell Growth Medium 2 (EGM2, Promocell, C-22111) supplemented with 10% Fetal Bovine Serum (FBS). Cell proliferation (Day 1-3) was monitored in multifrequency (MFT) mode for three days until the impedance reached a plateau. For endothelial barrier permeability studies (Day 4-5), cells were then transitioned to 0.1% FBS-supplemented EGM2 for 5 hours for baseline and challenged with 0.1 U/ml Thrombin (Sigma-Aldrich, T8885), with data collected overnight in single-frequency (SFT) mode at 4000 Hz. For migration studies (Day 5-6), cells were supplied with fresh 0.1% FBS EGM2 media, and a wound was created using the following setting: 60,000 Hz for 20 seconds. Migration was recorded in MFT mode overnight. Cell impedance, capacitance, and resistance signals collected through the ECIS assay were used for downstream analysis.

### Genome wide association between function and genotype

ECIS data were exported as.csv files. For the analysis we considered only 92 of the 100 individuals, removing the ones for whom errors occurred during the measurement. After initial quality control proliferation data was excluded due to the large batch effects and noise observed among the control group. For the remaining analyses we focused on wounding and thrombin challenge data using only the 4000 Hz frequency, as all collected frequencies were highly correlated (r2 > 0.97). Due to the inherent noise in the resulting functions, caused by measurements taken at closely spaced time points, the initial step involved smoothing the measurements using cubic B-splines (Ramsay 2005), with the smoothing parameter (λ) set to 1 and 1e-4 for thrombin and wounding, respectively. To allow for comparison, each of the thrombin functions (impedance, capacitance, resistance) were aligned, ensuring that peaks occurred at the same time point. To address batch effects in the data, which became evident when comparing the controls, each assay-measurement pair underwent a process of centering the batches with respect to their mean. This entailed removing the mean function for each batch, resulting in all batches having a zero mean. Subsequently, the mean function of the individual duplicates was computed to obtain a single function for each individual. For each pair of assay-measurement, a Functional Principal Component Analysis (FPCA) (Ramsay 2005) was performed across all functions, retaining only the first Functional Principal Component (FPC). Based on the FPCA results, we computed scores representing the projection of the functions onto the FPC. These scores enabled the ranking of the functions, and consequently of the samples, based on their overall behavior. These were then used to perform GWAS analysis using Plink v2.00a4.3 (Chang et al. 2015) (--glm option), with sex and the first two principal components (PCs) of the genetic relatedness matrix as covariates.

### Colocalization analyses

First of all, we defined the boundaries of significant genomic regions of association by identifying all the SNPs with a p-value lower than 1 × 10−5 for diseases, and 1 × 10−4 for HUVEC molecular QTLs. We then calculated the distance among each consecutive SNP below the set thresholds in the same chromosome: if two SNPs were less than 250 kb apart, they were assigned to same locus; otherwise, they were assigned to two different loci. We then considered as significant all the loci having a sentinel SNP with p-value < 5 × 10−8 for diseases, and < 2 × 10−5 for HUVEC molecular QTLs, and finally slightly enlarged these regions by adding ±100 kb.

After defining the genomic regions having a significant trait association, we performed conditional analysis using the GCTA-COJO v.1.91.4beta software (Yang et al. 2012). We employed the genotypes of 30,000 unrelated individuals of white British ancestry from UK Biobank (Bycroft et al. 2018) as LD reference panel and used DENTIST (Chen et al. 2021) prior to COJO to detect and eliminate variants showing heterogeneity between the GWAS and LD reference samples. First, we applied a stepwise forward conditional regression procedure (‘cojo-slct’) to select the independently associated SNPs at each trait-locus, among SNPs having MAF > 1% for diseases and 5% for molecular QTLs. The stopping criterion employed was for the conditional and joint p-values (pJ) to be larger than 1 × 10−4. This resulted in the identification of conditionally independent SNPs within the genomic region boundary. Then, for each of the identified variants, association analysis was re-performed conditional on all the other independent signals at the same locus (‘cojo-cond’), except for the target one (leave one out approach), generating conditional datasets each having only one independent association signal. For all conditional datasets containing at least a SNP with conditional p-value (pC) lower than 1 × 10−6 for diseases and 2 × 10−5 for molecular QTLs, loci boundaries were redefined using the same procedure previously described (using a SNPs inclusion p-value threshold of 1 × 10−4). Conditional datasets not satisfying this p-value criterion were discarded from further analyses. This process was iterated for all diseases and molecular QTLs exhibiting a significant association in the previously identified loci, obtaining a set of conditional datasets where the association signal is driven by a single causal variant.

We then fine-mapped each conditional dataset obtained by computing the 99% credible set, using the ‘finemap.abf’ function from the R package ‘coloc’ v.5.2.3 (Giambartolomei et al. 2014). We proceeded by pairwise testing for colocalisation, as implemented by Gianbartolomei et al. (Giambartolomei et al. 2014), only those conditional datasets sharing at least a SNP in their 99% credible set, using the ‘coloc.abf’ function from the R package ‘coloc’ v.5.2.3. We considered robust colocalisation tests resulting in posterior probability (PP) >= 0.75 for either H4 (hypothesis of same underlying causal variant contributing to both traits variation) or H3 (hypothesis of two independent underlying causal variants contributing to traits variation). Since ‘coloc’ package allows only for pairwise testing, we followed-up colocalisation analysis by network analysis (using the R package ‘igraph’ v.2.0.2) to identify, among multiple pairs of colocalising conditional datasets (PP.H4 >= 0.75), larger regulatory modules of traits all sharing the same causal variant.

### Data sources

GWAS summary statistics (Supplementary Table 2) were obtained from the following studies: stroke, GIGASTROKE Consortium meta analysis (Mishra et al. 2022); coronary artery disease (CAD), The CARDIoGRAMplusC4D Consortium (Aragam et al. 2022); Type 2 Diabetes (T2D), DIAMENTES database (Mahajan et al. 2022); pulmonary artery hypertension (PAH) from Rodhes et al. (Rhodes et al. 2019); atrial fibrillation (AF) from Nielsen et al. (Nielsen et al. 2018); cardiovascular disease (CVD) and venous thromboembolism (VTE) from Dönertaş et al. (Dönertaş et al. 2021); haemorrhoidal disease (HD) from Zheng et al. (Zheng et al. 2021); pulmonary heart disease and phlebitis and thrombophlebitis from Lee Lab (https://www.leelabsg.org/resources). Blueprint data was obtained from Kundu et al; (Kundu et al. 2022). Before performing the colocalization analysis the following quality control steps were applied: genome coordinates were converted to hg38 build with CrossMap (Zhao et al. 2014) and SNP markers names were updated with the new coordinates. Variance, z-scores and minor allele frequencies were calculated in R v4.1.0 (R Core Team, 2021). The number of causal SNPs was set to 2 and colocalization events with > 0.05 CLPP (colocalization posterior probability) were considered significant. Summary statistics for HUVEC - eQTL, sQTL, caQTL, hQTL generated in this study was used for the colocalization analyses.

Principal Component Analysis (PCA) of gene expression in endothelial cells from different sources.

Gene expression count data were obtained from ARCHS4 (https://maayanlab.cloud/archs4) (archs4_gene_human_v2.2.h5; Accessed 12/04/2023) (Lachmann et al. 2018) and combined with our samples gene expression counts. Details on the GEO series for each dataset are available (Suppl. Table S2). Batch effects were corrected using the ‘ComBat’ function from the ‘sva’ package (Leek et al. 2012), specifying two batches: one for our data and another for all others. Our HUVEC and HUVEC samples obtained from ARCHS4 were treated as the same biological type. PCA plots were generated using ggplot2.

